# Neural representations of beliefs in a multi-dimensional inference task

**DOI:** 10.64898/2025.12.31.697231

**Authors:** Patrick Q. Zhang, Michael J. Jutras, Adam J.O. Dede, Edgar Y. Walker, Elizabeth A. Buffalo, Adrienne L. Fairhall

## Abstract

Adaptive behavior requires maintaining and updating probabilistic beliefs about the world, yet how distributed brain circuits implement such computations remains unknown. We recorded from over 1,400 neurons across six brain regions in monkeys performing a multi-dimensional inference task requiring them to infer hidden rules through trial-and-error learning. Behavior was well-described by models based on Bayesian updating of beliefs over rule features. Neural representations of both observable variables (stimuli, rewards) and latent beliefs (rule preferences, confidence) were broadly distributed across hippocampus, amygdala, prefrontal cortex, anterior cingulate, striatum, and inferior temporal cortex. Belief representations were present throughout all task periods but exhibited region- and epoch-specific dynamics. Critically, trial-to-trial changes in population activity reflected Bayesian belief updating: neural responses evolved according to the integration of prior beliefs with new evidence. Additionally, we identified confidence representations that were independent of specific beliefs and showed distinct temporal profiles. These results demonstrate that probabilistic inference emerges from coordinated dynamics across distributed brain systems, with different regions contributing flexibly according to computational demands at different states of learning and decision-making.

## 1 Introduction

To navigate the world effectively, organisms must distill high-dimensional sensory input into structured representations that support goal-directed behavior. A central challenge in understanding cognition is explaining how the brain transforms continuous experience into an organized model of the world that can guide current and future decisions. Here, we investigate how such an internal model may be implemented in the primate brain using a multi-dimensional inference task that requires animals to integrate observations over time to guide behavior. We formalize this problem using a Bayesian framework, which allows us to precisely quantify what animals should represent—beliefs about hidden states of the world—and how those representations should change through Bayesian inference following new observations. This approach provides specific predictions about the computations the brain must perform to solve the task. To investigate the neural basis of these computations, we performed large-scale recordings focused on six interconnected brain systems previously implicated in learning and sequential decision-making: value-based decision circuits (dorsal striatum, amygdala) [1–8]; cognitive control regions (lateral prefrontal cortex and anterior cingulate cortex) [9–14]; memory circuits (hippocampal formation) [15–18]; and high-level sensory cortex (inferior temporal cortex) [19–22]. We find that representations of belief and other task variables are broadly distributed across all recorded regions rather than localized to specific circuits, with both stable and dynamic encoding observed across different trial epochs. Critically, neural activity evolves trial-by-trial in a manner consistent with Bayesian updating: population responses reflect how new observations change beliefs according to the inference model. Moreover, belief representations exhibit a specific geometric organization in that they are non-orthogonal to one another and structured according to the visual feature dimensions that define the task space. Together, these findings reveal how multiple brain systems coordinate to maintain and update probabilistic beliefs during naturalistic inference.

## 2 Results

### 2.1 Monkeys perform a complex multi-dimensional inference task

To study the neural representation of beliefs, we trained monkeys on a multi-dimensional inference task [23], in which they must infer a hidden rule governing reward through trial and error. On each trial, monkeys search a visual scene consisting of four stimuli, each defined by a unique combination of color, shape, and pattern. They select a stimulus by making a saccade to it and maintaining fixation on the selected stimulus for 800ms. At this point, all stimuli disappear from the screen and feedback is delivered: food slurry reward for a correct choice or a timeout for an incorrect choice. Feedback is determined by whether the chosen stimulus contains a feature specified by the hidden rule (e.g., ‘green’ or ‘striped’). After consistent correct responses (8 out of 8, or 16 out of the 20 past trials), the hidden rule switches, uncued, to a different feature chosen at random, and the monkey has to infer the new hidden rule via trial and error (Fig 1a,b). Critically, each stimulus contains three features (one color, one shape, and one pattern), so a single trial’s feedback is ambiguous - it does not reveal which feature dimension defines the rule. Thus, to maximize reward, subjects must integrate a history of stimulus selections and feedback from multiple past trials to guide future decision making.

**Fig. 1:**
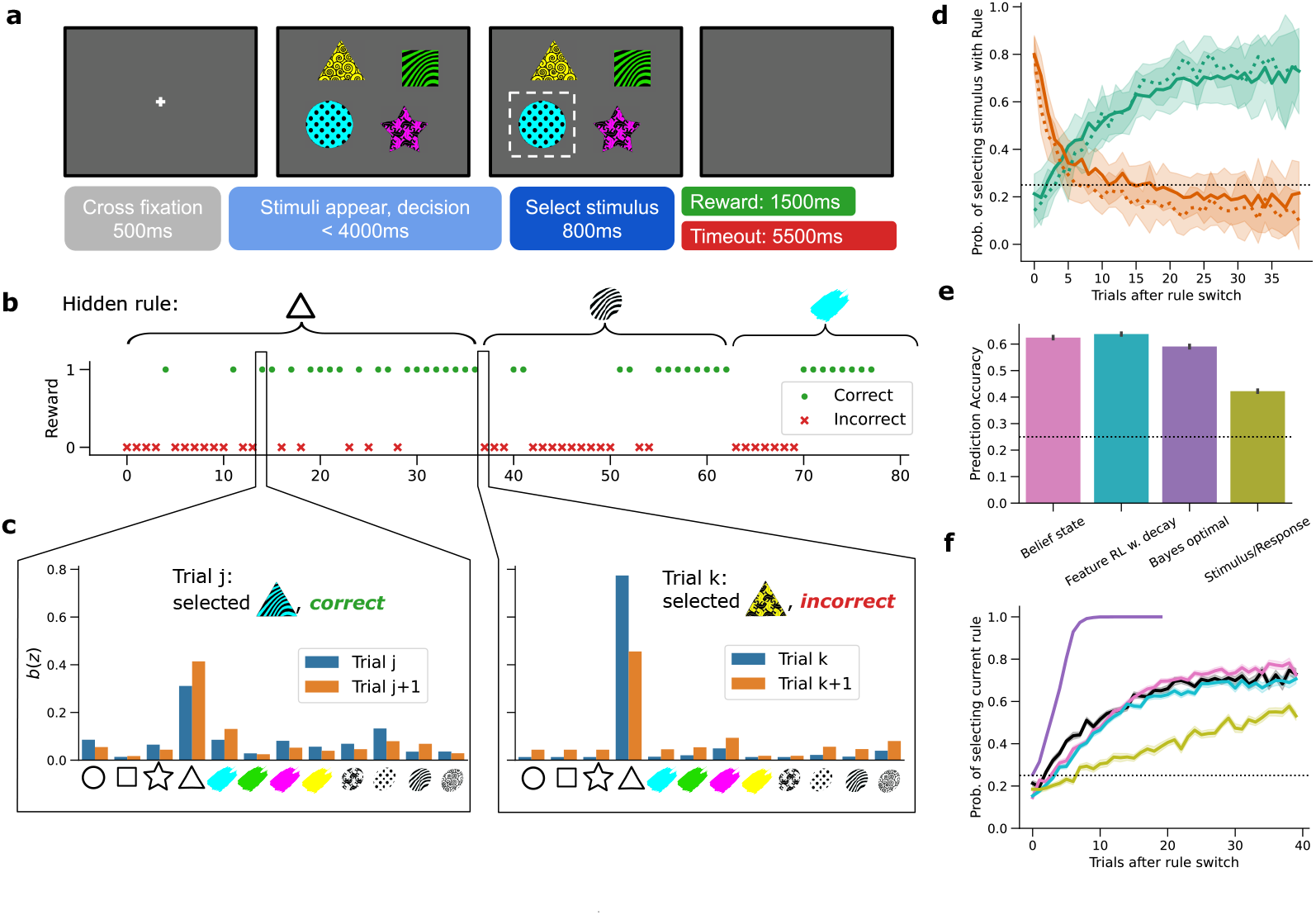
Monkeys perform inference on a multi-dimensional choice task. **a**, Trial structure. Monkeys initiate trials by fixating on a central cross. Then, four stimuli appear, each composed of three features: a color, shape, and pattern. Monkeys select a stimulus by maintaining fixation on it (dotted white boundaries), and receive reward if the selected stimulus contains the hidden rule feature (1 of 12 possible features). **b**, Session structure. The hidden rule remains consistent within blocks of trials and switches uncued after monkeys reach a performance criterion (vertical dotted lines). **c**, Example belief state distributions inferred from the behavioral model. Beliefs are updated recursively on each trial based on selected stimulus features and trial outcome. **d**, Performance curves for Monkey S following rule switches. The probability of selecting a stimulus containing the previous rule (orange) decreases while the probability of selecting a stimulus containing the current rule (green) increases. Solid lines indicate monkey behavior, dotted lines indicate Belief state model predictions. Shaded areas represent standard deviation across rule switches. **e**, Prediction accuracies for Monkey S’s stimulus selections for each behavioral model, evaluated on held-out test sessions. **f**, Probability of selecting stimuli containing the current rule after rule switches for Monkey S (black) and different behavioral models. Shaded areas represent s.e.m. across rule switches

The animals successfully learned this task. Following rule switches, monkeys infer the new rule within tens of trials, as evidenced by their gradually increasing reward rates. (Fig 1d, solid green lines). Initially, monkeys tend to perseverate on the previously rewarded feature for several trials before switching their strategy (Fig 1d, solid orange lines).

These behavioral patterns suggest that monkeys decompose the visual stimuli into constituent features, and integrate information across trials to guide inference. To test this hypothesis quantitatively and to estimate the animals’ internal beliefs, we compared several computational models of task performance (Figs 1e,f, S1). We compared four candidate strategies. If monkeys do not take advantage of the feature-based structure of the task, they must rely on stimulus-response memorization, learning the correct response for each of the many possible stimuli (Stimulus/Response, olive). Alternatively, monkeys could exploit feature structure using three different approaches: (1) Bayesian inference with known task parameters (Bayes Optimal, purple), which computes posterior beliefs over rule features; (2) feature-based reinforcement learning (Feature RL w. decay, cyan), which incrementally updates values assigned to individual stimulus features, assuming a finite memory window; and (3) approximate Bayesian inference (Belief state, pink), which updates beliefs defined over features but assumes imperfect knowledge of task statistics. We fit free parameters in the feature RL with decay and belief state models using maximum likelihood estimation on monkeys’ trial-by-trial choices (Methods 5.4.5).

We estimated each model’s performance on the actual stimulus sequences experienced by the monkeys and compared the resulting learning curves to observed behavior (Figs 1f, S1c,d,g,h). Monkeys (black) fell short of the Bayes optimal model (purple) but performed substantially better than the stimulus-response memorization strategy (olive), indicating their use of a feature-based strategy. Both the feature RL with decay model (cyan) and the belief state model (pink) captured behavioral performance approximately equally well (Figs 1e, S1a,b), suggesting the monkeys decompose stimuli into features and track the reward associations of individual features over time.

Although these two models differ mechanistically, both track estimates of each feature’s importance through trial-by-trial updates. In the RL model, each feature is assigned a value that increases when rewarded and decreases through temporal decay, capturing finite memory. On each trial, feature values within each stimulus are summed, and stimulus selections are made probabilistically according to total stimulus values (Methods 5.4.3). By contrast, in the Bayesian belief state model, subjects maintain a probability distribution over which feature is currently the rule. During stimulus selection, monkeys apply these beliefs to estimate which stimulus is most likely to contain the hidden rule feature. After feedback, monkeys update their beliefs by integrating the prior distribution with information from the current trial (selected features and outcome) according to an assumed internal model of the task structure [24, 25] (Methods 5.4.2). Because monkey behavior is not Bayes optimal (Figs 1f, S1c,d,g,h), here we have assumed that the monkeys’ internal model of the task environment differs from the true task structure [25]. Specifically, we assume that monkeys treats rewards as probabilistic rather than deterministic, which accounts for their suboptimal but robust performance.

To link behavior to neural activity, we needed to estimate the monkeys’ trial-by-trial beliefs about the hidden rule. Both the RL and Bayesian belief state models provide such estimates through internal variables - feature values and belief distributions, respectively - that are highly correlated with one another (Fig S1b,f). We selected the Bayesian belief state model for subsequent neural analyses because it represents beliefs as an explicit probability distribution over the possible rule features, and accurately captures behavioral patterns including perseveration dynamics (Figs S1d,h). This framework allowed us to ask how neural populations encode and update probabilistic beliefs during multi-dimensional inference.

### 2.2 Distributed representations of stimulus features and reward

To identify how the primate brain performs this task, we recorded from 1422 single units in multiple brain regions (1089/333 units in Monkey S, B respectively), Table 1, Fig S2). A substantial number of recorded units were located in brain areas previously implicated in various aspects of sequential decision making, learning, and memory: the lateral prefrontal cortex (LFPC, 239 units), anterior cingulate cortex (ACC, 85 units), dorsal striatum (DSTR, 111 units), amygdala (AMY, 97 units), inferior temporal cortex (ITC, 241 units), and hippocampal formation (HC, 131 units).

**Table 1.** Number of units per region. Regions above the dashed line were additionally analyzed as individual populations.

| Region | Monkey S Units | Monkey B Units | Combined |
| --- | --- | --- | --- |
| all regions | 1089 | 333 | 1422 |
| inferior temporal cortex (ITC) | 185 | 56 | 241 |
| lateral prefrontal cortex (LPFC) | 239 | 0 | 239 |
| hippocampal formation (HC) | 48 | 83 | 131 |
| dorsal striatum (DSTR) | 77 | 34 | 111 |
| amygdala (AMY) | 73 | 24 | 97 |
| anterior cingulate cortex (ACC) | 85 | 0 | 85 |
| claustrum | 77 | 7 | 84 |
| superior parietal cortex | 61 | 16 | 77 |
| motor cortex | 63 | 0 | 63 |
| posteromedial cortex | 42 | 15 | 57 |
| inferior parietal cortex | 43 | 11 | 54 |
| thalamus | 33 | 16 | 49 |
| orbitofrontal cortex | 35 | 0 | 35 |
| extrastriate visual areas V2–V4 | 8 | 25 | 33 |
| primary visual cortex | 12 | 16 | 28 |
| perirhinal/parahippocampal cortex | 1 | 23 | 24 |
| insular cortex | 7 | 7 | 14 |

According to the Bayesian belief state model, beliefs are updated each trial by integrating information about selected stimulus features and reward feedback (Fig 1 c). We therefore first asked how these externally observable task variables are represented in neural activity, at both single-unit and population levels.

In each region, we identified neurons whose firing rates were significantly modulated by selected stimulus features (e.g. whether a selected stimulus contained a specific shape such as a triangle). These neurons exhibited diverse tuning properties and temporal dynamics (Figs 2a, S3c). Across the entire population, we quantified both the fraction of units significantly encoding selected stimulus features and the fraction of variance in neural activity explained by these features. Both metrics peaked during fixation on the selected stimulus (Fig S3a). Notably, both metrics persisted into the feedback period, even after stimuli disappeared from the screen, indicating sustained representation of choice information. This pattern was observed across most regions, though encoding strength was notably weaker in the ACC (Fig S3 b).

**Fig. 2:**
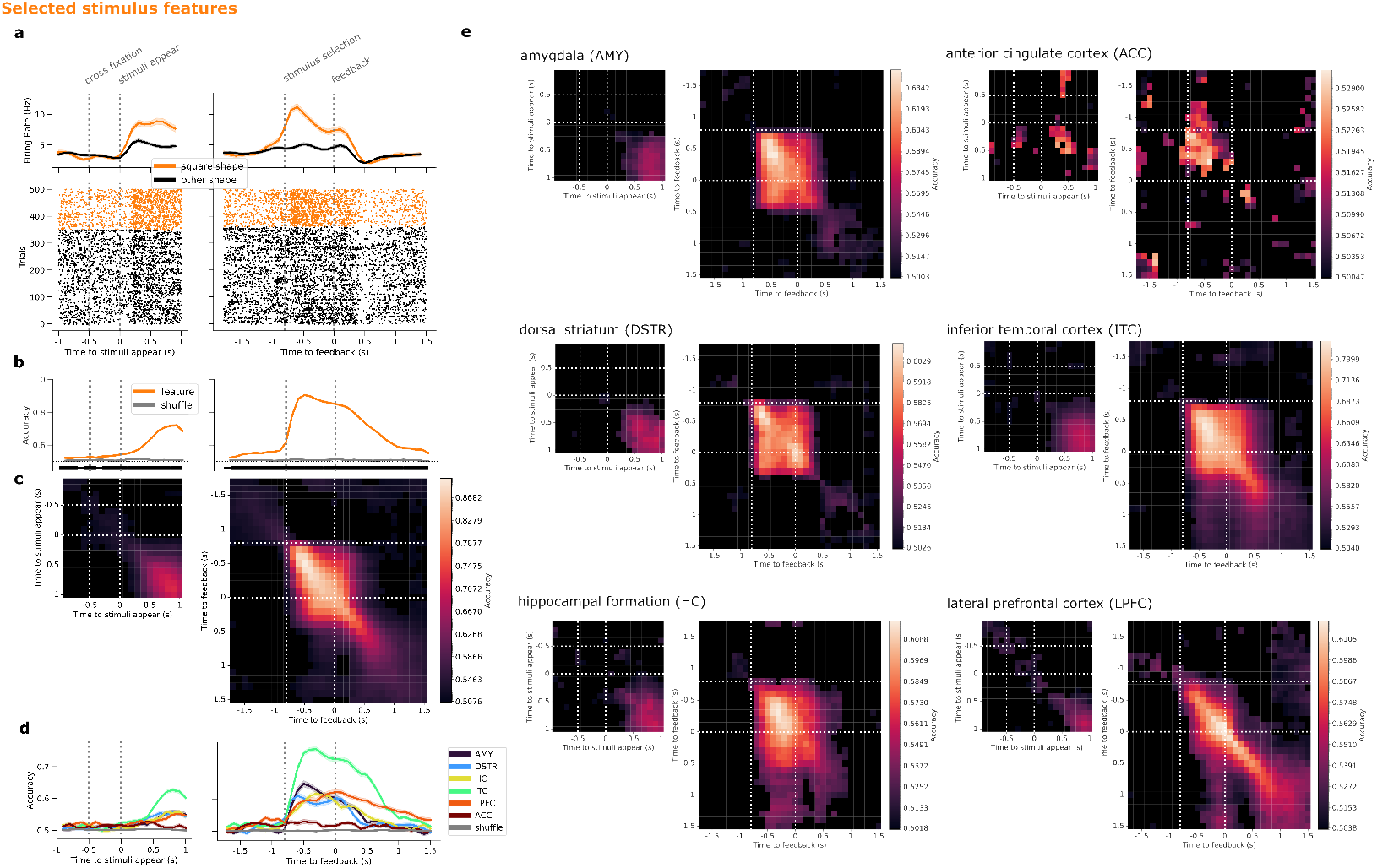
Neural representations of selected stimulus features across trial periods. Neural activity is aligned to stimulus onset and feedback onset. Dotted vertical lines indicate trial events. **a**, Sample ITC neuron selective for whether a square-shaped stimulus was selected. Top: firing rates averaged by condition, shaded regions represent s.e.m. across trials. Bottom: spike rasters from example trials, grouped by condition. **b**, Population decoding accuracy for selected stimulus features, averaged across all features and trained/tested in 100ms time bins. Horizontal black bar thickness indicates significance level: thick: *p* < 0.001, moderate: *p* < 0.01, thin: *p* < 0.05. **c**, Cross-temporal decoding accuracy for selected stimulus features. Decoders are trained in each time bin (rows), and evaluated in each time bin (columns). Black: non-significant bins (*p* ≥0.01). **d**, Population decoding accuracy for selected stimulus features, evaluated separately for each brain region. **d**, Cross-temporal decoding accuracy in the format of **c**, evaluated separately for each brain region

To examine when selected stimulus features are decodable from population activity, we trained linear classifiers on neural firing rates in non-overlapping 100ms time bins throughout each trial. Decoding accuracy peaked during fixation on the selected stimulus and persisted into the feedback period (Fig 2b), consistent with the single-unit encoding results.

To assess whether the population representation remains stable or transforms over time, we computed a *cross-decoding accuracy (CDA)* matrix: each row shows the decoding accuracy of a decoder trained on neural activity in time bin *i* (Fig 2c, rows) when evaluated on all other time bins *j* (Fig 2c, columns). High off-diagonal accuracy indicates stable representations across time, while high accuracy confined to the diagonal indicates time-varying, dynamic representations.

We observed a clear block structure in the CDA matrix: a stable representation of selected stimulus features during stimulus fixation that persisted 200–300ms into the feedback period, followed by a transformation to a different stable representation later in the feedback period (evident in the lower-right block). This suggests that the population stores the stimulus representation in a different basis during the feedback period. Critically, units that contribute to this population representation were not localized to any single region, but were distributed across all recorded areas (Fig S3d), suggesting that information about selected features is broadcast widely across decision-making circuits.

Examining decoding accuracy (Fig 2d) and corresponding CDA matrices computed separately for each region (Fig 2e) reveals distinct contributions to stimulus feature representation.

Inferior temporal cortex (ITC) showed the strongest feature decoding, as expected given its role in visual object processing [26–28]. Decoding in ITC peaked 300ms after stimulus fixation onset, with a stable representation during the visual period that then transformed to a distinct representation in the feedback period, when stimuli have disappeared from the screen. Stimulus identity is stably represented only during the selected stimulus fixation period in amgydala (AMY) and dorsal striatum (DSTR), and, with a slight delay, in hippocampus (HC), all with comparable decoding accuracy. Peak decoding accuracy in AMY and DSTR occurred around 300ms after stimulus fixation, similar to ITC. By contrast, lateral prefrontal cortex (LPFC) showed peak decoding accuracy near the end of the fixation period, suggesting delayed or more sustained processing; the representation in LPFC also shows a change, with distinct decoder weights (Fig S3d), during the feedback period. Both ITC and LPFC maintained distinct representations during the feedback period, indicated by the block structure in their CDA matrices. This sustained encoding after stimulus offset suggests that these regions maintain choice-related information during the period when beliefs are updated based on feedback. In contrast to other regions, the anterior cingulate cortex (ACC) showed minimal feature representation at any point in the trial, suggesting it may primarily encode other task variables (see below).

Reward outcome (correct vs. incorrect trials) was also broadly represented across brain regions. Single neurons showed diverse reward tuning across multiple areas (Figs 3a, S4c), and population-level decoding of feedback achieved near-perfect accuracy (> 95%) during the feedback period (Fig 3b).

**Fig. 3:**
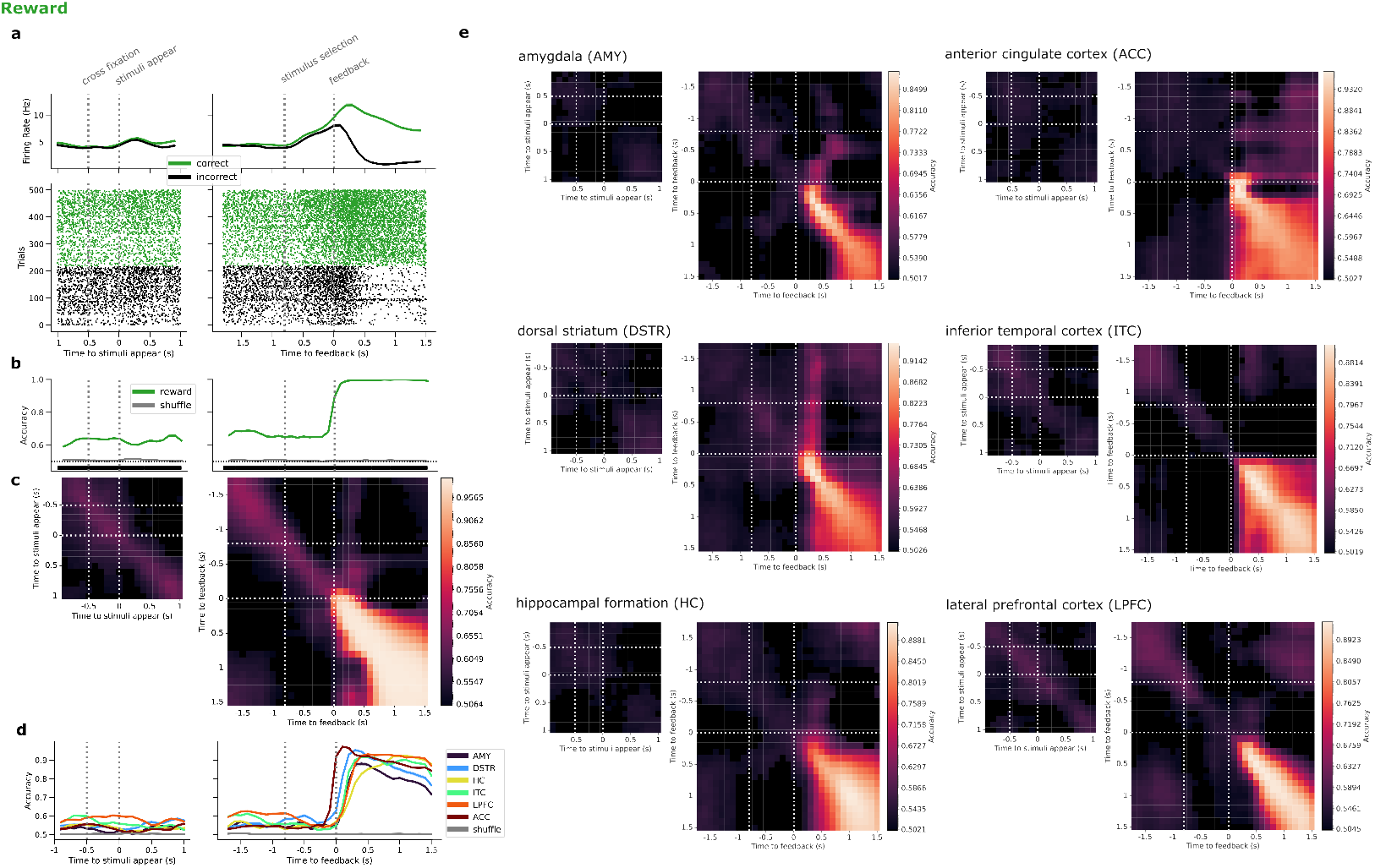
Neural representations of reward across trial periods. Neural activity is aligned to stimulus onset and feedback onset. Dotted vertical lines indicate trial events. **a**, Sample ACC neuron selective for trial outcome. Top: firing rates averaged by condition, shaded regions represent s.e.m. across trials. Bottom: spike rasters from example trials, grouped by condition. **b**, Population decoding accuracy for reward, trained/tested in 100ms time bins. Horizontal black bar thickness indicates significance significance level: thick: *p* < 0.001, moderate: *p* < 0.01, thin: *p* < 0.05. **c**, Cross-temporal decoding accuracy for reward. Decoders trained in each time bin (rows) and evaluated in each time bin (columns). Black: non-significant bins (*p*≥ 0.01). **d**, Population decoding accuracy for reward, evaluated separately for each brain region. **d**, Cross-temporal decoding accuracy in the format of **c**, evaluated separately for each brain region

Region-specific analyses revealed two key organizational principles. First, reward signals showed a striking temporal progression: significant decodability is present in ACC almost immediately after feedback onset, followed by DSTR, then the LPFC, ITC, and AMY, and finally HC, peaking almost 1s later (Fig 3d). This sequential activation may reflect a cascade from immediate outcome detection to reward-related memory consolidation.

Second, cross-temporal decoding revealed region-specific representational dynamics (Fig 3e): ACC and ITC maintained stable reward representations, particularly late in the feedback period, while LFPC showed more dynamic, time-varying representations. The stability of the representation in ACC and ITC suggests activation of a fixed neural population, while in LPFC, the dynamic representation suggests that there, trial outcome generates a unfolding pattern of neural dynamics. This may reflect ongoing planning or integration with other evolving task variables.

### 2.3 Dynamical representations of beliefs throughout trial periods

Beliefs should play distinct roles throughout the task. During visual search and stimulus selection, beliefs influence one’s search strategy and inform which stimuli likely to yield reward. During feedback, beliefs must be integrated with outcome information in order to be updated for the next trial. Between trials, beliefs must be maintained to inform future decisions. This functional diversity suggests that belief representations should be present throughout all trial periods, but may be organized differently across brain regions and task epochs according to computational demands. We therefore asked whether and how the beliefs predicted by our behavioral model are represented in neural activity across time and brain regions.

Identifying neural signatures of belief in this task presents two challenges. First, with 12 possible rule features, the full belief state is a 12-dimensional probability distribution that cannot be directly decoded given our sample sizes. We therefore probe beliefs one feature at a time, asking how strongly the network represents belief about each individual feature. Second, beliefs are inherently correlated with observable task variables. For example, trials with high belief that “triangle” is the rule also tend to be trials in which the probability of selecting a triangle stimulus is high, making it difficult to distinguish neural signals of believing triangle is correct from signals related to seeing and selecting a triangle (Fig 2). Similarly, high-confidence beliefs often occur on rewarded trials, confounding belief strength (confidence) with reward signals during the feedback period (Fig 3).

Given the high dimensionality of the problem, we developed a partitioning approach that isolates belief-related neural signals for each feature. For each feature *X*, we categorize all trials into three belief states based on the model’s predicted posterior distribution: low confidence, when belief is distributed relatively uniformly across features (Fig 4, left), high confidence in *X*, when belief strongly favors feature *X* (Fig 4, top right), and high confidence in other, when belief strongly favors some feature other than *X* (Fig 4, bottom right). These three partitions define two orthogonal axes for probing belief representations. The preference axis (Fig 4, blue), contrasts high confidence in *X* with high confidence in other features, revealing whether neural activity distinguishes which feature is believed to be the rule. The confidence axis (Fig 4, red) contrasts low versus high confidence states, revealing whether neural activity reflects how certain the monkey is, independent of which feature is preferred.

**Fig. 4:**
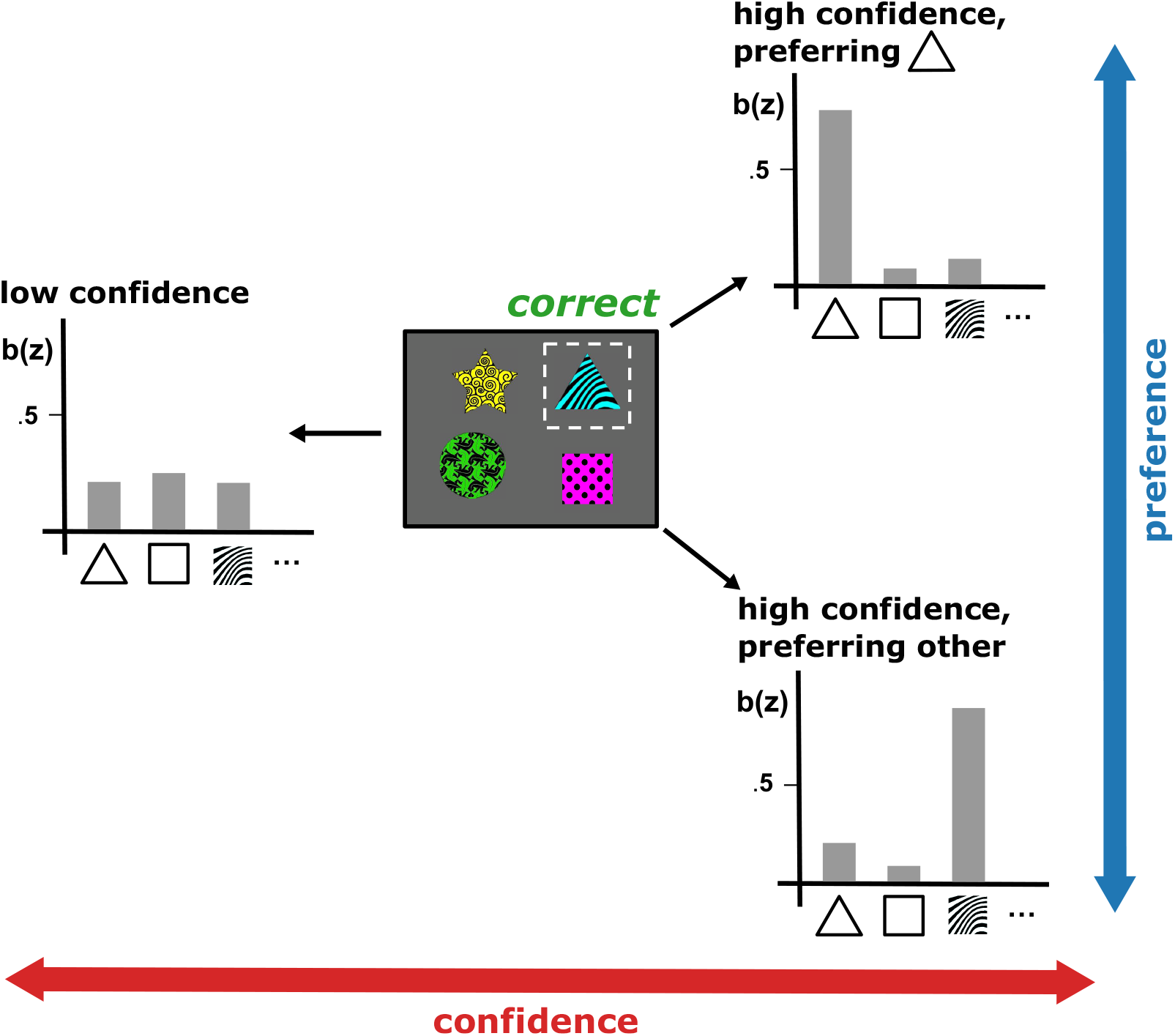
Partitioning belief states to isolate confidence and preference representations. An observed stimulus selection and reward outcome (e.g. selecting a stimulus containing a triangle and receiving reward; middle) can correspond to multiple hidden belief states. For each feature (e.g. triangle), we partition inferred belief states into three categories: low confidence (left) where beliefs are distributed relatively uniformly across all features; high confidence in triangle (top right) where belief strongly favors triangle as the rule, and high confidence preferring other (bottom right) where belief strongly favors a feature other than triangle. These three partitions define two orthogonal axes for probing belief representations: confidence (red), which contrasts low versus high confidence states, and preference (blue), which contrasts high confidence in one feature versus high confidence in another feature.

Critically, to eliminate confounds from task variables, we restrict our analysis to trials with identical stimulus selections and reward outcomes: trials in which the monkey selected a stimulus containing feature *X* and received reward (Fig 4, middle). Within this matched set of trials, any differences in neural activity across belief partitions must reflect the latent belief state rather than sensory or reward signals. This approach allows us to ask how neural populations encode both belief preference and confidence throughout the trial, independent of confounding observable variables (Methods 5.5).

Using this approach, we identified neurons in all regions whose firing rates were significantly modulated by belief preference - that is, by which feature the monkey believed to be the rule. Single units selective for preference were observed across all regions and throughout all trial periods (Figs 5a, S5c). At the population level, both the fraction of units encoding preference and the fraction of variance explained by preference increased across the trial, peaking during the feedback period (Fig S5a). Population-level decoding revealed that preference was significantly decodable throughout all trial periods (Fig 5b). However, cross-temporal decoding showed that preference representations were more dynamic than those of observable task variables (selected feature and reward), with rapid transformations across trial epochs, though some stability was evident during the visual search prior to choice (Fig 5c).

**Fig. 5:**
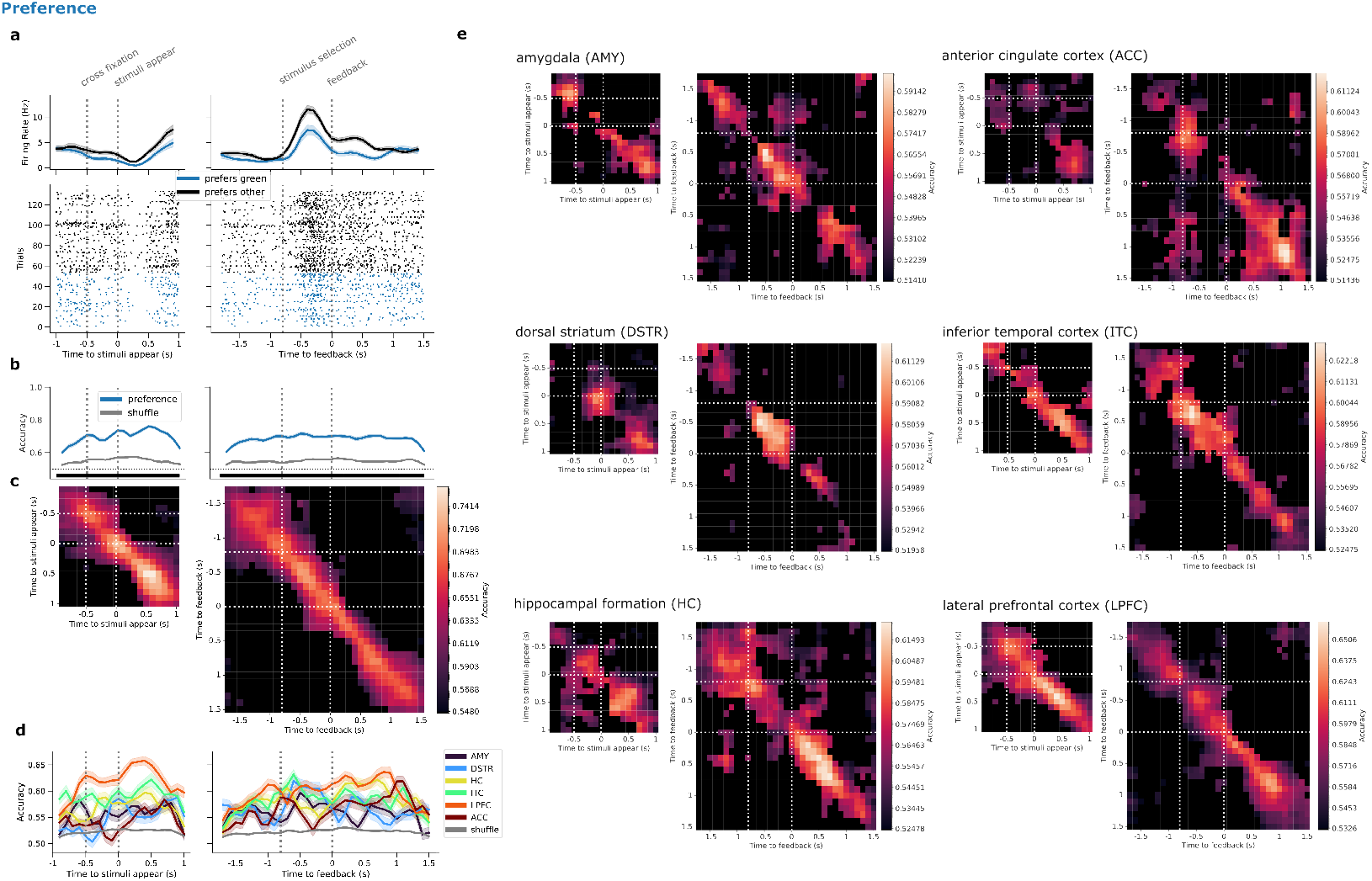
Neural representations of belief preference across trial periods. Neural activity is aligned to stimulus onset and feedback onset. Dotted vertical lines indicate trial events. **a**, Sample LPFC neuron selective for belief prefence: whether the monkey believed green was the rule versus favored another feature, as predicted by the belief state model. Analysis was restricted to trials in which a green stimuli was selected and rewarded. Top: firing rates averaged by condition, shaded regions represent s.e.m. across trials. Bottom: spike rasters from example trials, grouped by condition. **b**, Population decoding accuracy for preference, averaged across all features and trained/tested in 100ms time bins. Horizontal black bar thickness indicates significance significance level: thick: *p* < 0.001, moderate: *p* < 0.01, thin: *p* < 0.05. **c**, Cross-temporal decoding accuracy for preference. Decoders trained in each time bin (rows), and evaluated in each time bin (columns). Black: non-significant bins (*p* ≥ 0.01). **d**, Population decoding accuracy for preference, evaluated separately for each brain region. **d**, Cross_temporal decoding accuracy in the format of **c**, evaluated separately for each brain region.

Region-specific analyses revealed distinct temporal profiles of preference encoding (Fig 5d). LPFC showed the strongest and most sustained preference representation, peaking prior to the appearance of the stimulus and remaining elevated throughout the trial. While the LPFC representations were generally dynamic, they exhibited relative stability in the pre-stimulus period, potentially reflecting advance preparation, or “holding in mind”, of rule-related representations. IT, AMY and HC all showed time-varying, decodable preference representations throughout the trial; notably, HC showed relatively stable encoding during the pre-selection period. By contrast, DSTR encoded preference primarily during stimulus presentation, while ACC showed preference encoding predominantly during the feedback period (Fig 5e).

Complementing preference representations, we also identified neurons in all regions whose firing rates were significantly modulated by belief confidence–how certain the monkey is about its belief–with modulation present throughout all trial periods (Figs 6a, S6c). In contrast to preference, which peaked during feedback, population-level confidence encoding showed a bimodal temporal profile, peaking first in the pre-stimulus period and again during feedback (Figs 6b, S6a). Cross-temporal decoding revealed multiple stable representational epochs, evident as distinct blocks in the CDA matrix: (1) pre-stimulus preparation, (2) visual search, (3) immediately post-feedback, and (4) late feedback period (Fig 6e, colored blocks). Representations of confidence transformed between these stable epochs, shown by low cross-decoding between blocks (Fig 6e, dark diagonals).

**Fig. 6:**
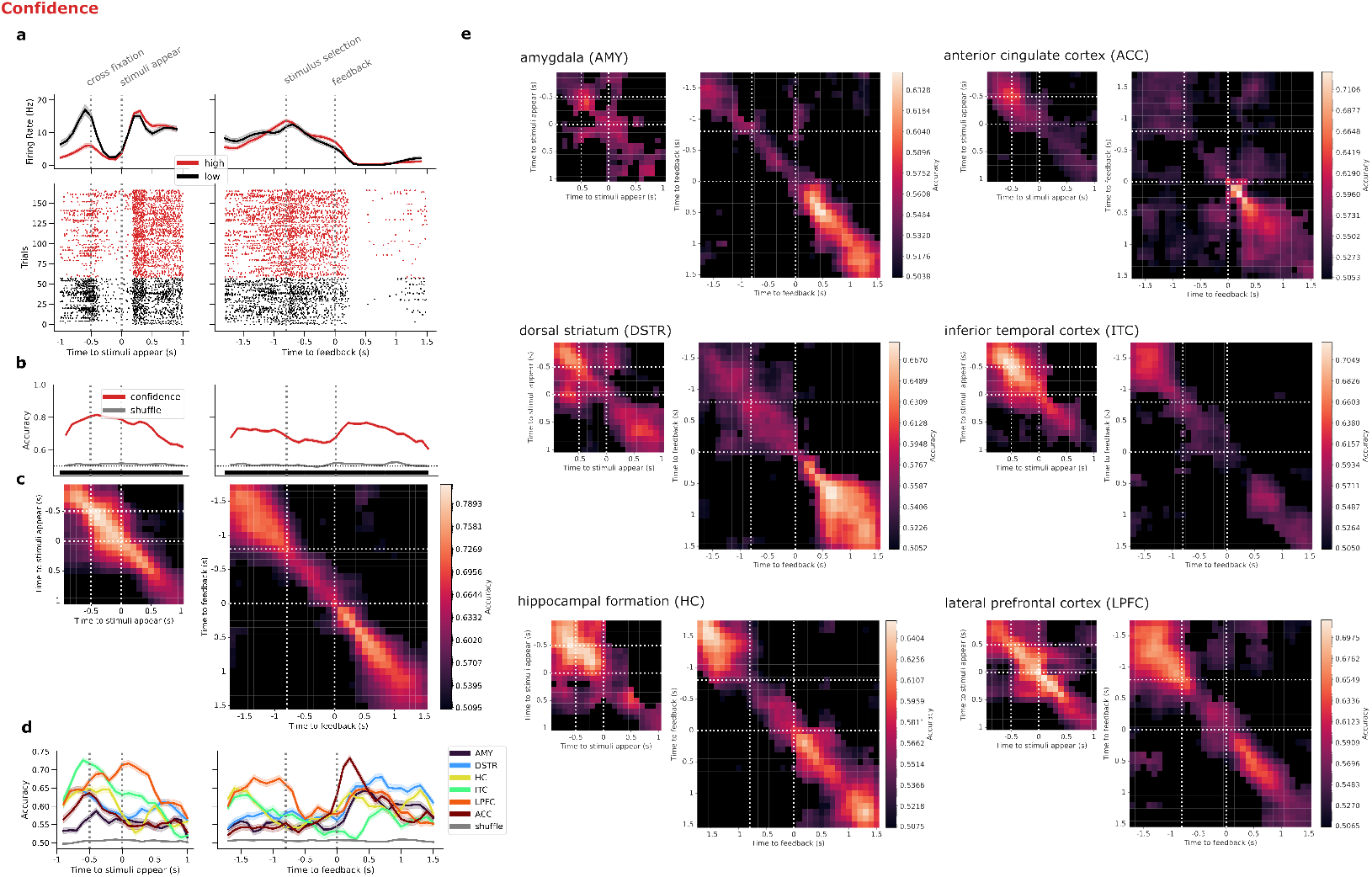
Neural representations of belief confidence across trial periods. Neural activity is aligned to stimuli onset and feedback onset. Dotted vertical lines indicate trial events. **a**, Sample ITC neuron selective for belief confidence: whether the monkey had low or high confidence in its belief about the hidden rule, as predicted by the belief state model. Analysis restricted to rewarded trials. Top: firing rates averaged by condition, shaded regions represent s.e.m. across trials. Bottom: spike rasters from example trials, grouped by condition. **b**, Population decoding accuracy for confidence, trained/tested in 100ms time bins. Horizontal black bar thickness indicates significance significance level: thick: *p* < 0.001, moderate: *p* < 0.01, thick: *p* < 0.05. **c**, Cross-temporal decoding accuracy for confidence. Decoders trained in each time bin (rows), and evaluated in each time bin (columns). Black: non-significant bins (*p*≥0.01). **d**, Population decoding accuracy for confidence, evaluated separately for each brain region. **d**, Cross-temporal decoding accuracy in the format of **c**, evaluated separately for each brain region.

Region-specific analyses revealed that different brain areas contributed to confidence during specific trial epochs (Fig 6d). Notably, confidence was weakly represented across all regions during the stimulus selection fixation period. ACC and DSTR both showed strongest confidence encoding during feedback, with ACC exhibiting a particularly strong but brief response immediately post-choice that then transitioned to more sustained encoding throughout the feedback period. Both regions also showed weaker confidence representations during the pre-stimulus interval. LPFC encoded confidence strongly during both the pre-trial and decision-making periods, maintaining representations across these epochs. HC showed a similar pattern to LPFC – strong and stable during pre-trial and decision-making periods, followed by strong but dynamic encoding during feedback – though with lower overall decoding accuracy. ITC showed only weak confidence encoding during search, choice and feedback, but exhibited stable representation during pre-stimulus period, potentially reflecting preparation for upcoming visual processing demands (Fig 6e).

### 2.4 Neural belief representations update according to Bayesian inference

In the Bayesian belief state model, beliefs are updated each trial through Bayesian inference, integrating prior beliefs with new observations. Critically, both the direction and magnitude of updates depend on the trial outcome and the prior belief state. For example, belief in feature *X* changes little when averaged across all correct or incorrect trials, but shows much larger changes when the selected stimulus contains feature *X* (7a, top). Moreover, the magnitude of change depends on prior confidence: belief in *X* increases most when confidence is initially low and a stimulus containing feature *X* yields reward but decreases most when confidence in *X* is initially high, yet selecting a stimulus containing *X* yields no reward (7a, bottom).

We asked whether neural population activity evolves across consecutive trials in a manner consistent with these predicted belief updates. We focused on the inter-trial interval - specifically, the 1-second period before stimuli appear, while the monkey maintained central fixation - reasoning that during this period the updated belief state is carried forward to guide the next trial. We computed the trial-to-trial change in population activity during this interval and grouped trials by the previous trial’s stimulus selection (selected *X* or not), outcome (correct/incorrect) and prior belief state (low confidence, high confidence in *X*, or high confidence in other). We then projected these activity changes onto the decoder axes for preference and confidence (computed without conditioning on selected stimulus features or reward) to measure how belief-related activity evolves.

We found that receiving correct feedback resulted in positive changes in confidence-related activity, while incorrect feedback resulted in negative changes (Fig 7b, top). This pattern held regardless of which features were selected, confirming that our confidence representation is feature-independent. Furthermore, when conditioned on prior belief states, the magnitude of updates matched model predictions: correct trials produced the strongest positive update when prior confidence was low, while incorrect trials produced the strongest negative update when prior confidence was high (7b, bottom).

**Fig. 7:**
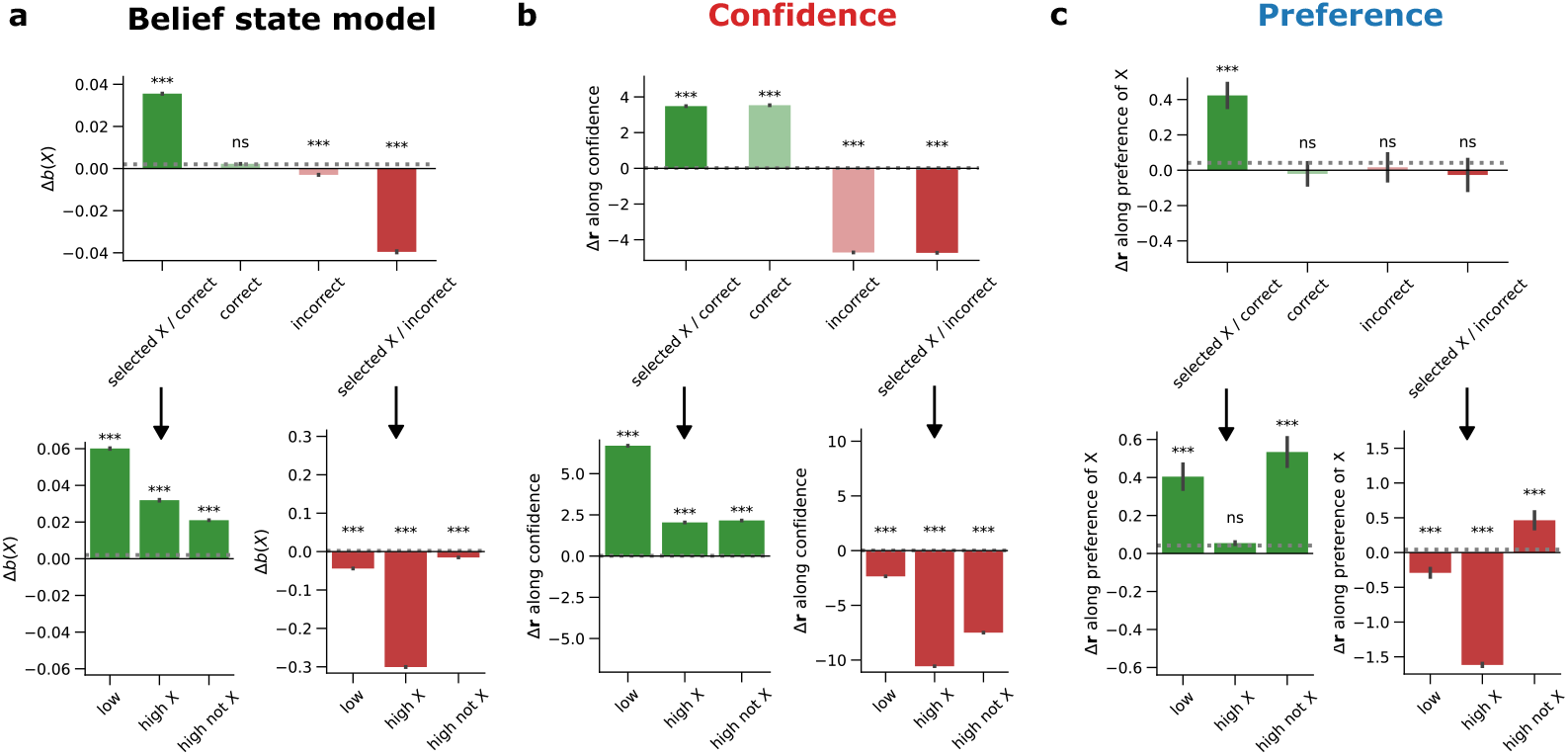
Trial-by-trial changes in belief-representing neural activity align with Bayesian inference. **a** Predictions from the Bayesian belief state model for how belief in feature *X* changes based on trial outcome (top) and prior belief state (bottom). **b** Change in neural population activity from the current trial to the next trial, measured during the 1s pre-stimulus interval and projected onto the confidence decoder axes. Changes are grouped by trial outcome (top) and prior belief states (bottom). **c** Same as **b**, but projected onto the preference decoder axes. Stars indicate significance levels relative to corresponding shuffles: ***: *p* < 0.001, **: *p* < 0.01, *: *p* < 0.05

For preference toward feature *X*, however, only one condition produced significant positive change along its corresponding preference axis: when feature *X* was selected and rewarded (Fig 7c, top). When conditioned on prior belief states, preference-related activity updates qualitatively matched model predictions in two of the three conditions: low confidence and high confidence favoring *X* (Fig 7c, bottom). However, when prior belief strongly favored a feature other than *X*, selecting a stimulus with *X* produced a positive change in *X*-preference activity, regardless of feedback outcome - deviating from model predictions. This deviation may reflect behavioral biases such as perseveration on recently selected features that are not fully captured by the normative Bayesian model.

### 2.5 Belief representations are aligned in neural activity space

Having identified neural representations of belief preference for individual features, we next asked how these representations are organized geometrically across the population. One hypothesis is that beliefs about different features are encoded orthogonally, such that neural activity associated with increasing belief in one feature is uncorrelated with activity associated with increasing belief in another feature. A toy example of this organization is shown in Fig. 8a, where individual neurons separately encode beliefs in different features through their firing rates, resulting in orthogonal coding dimensions in population activity space. Alternatively, encoding directions for different feature beliefs might be partially aligned, such that increasing belief in one feature drives population activity in a shared direction as increasing belief in another feature. In a simple example, adding a feature-agnostic neuron that encodes only confidence would produce such alignment (Fig. 8c).

**Fig. 8:**
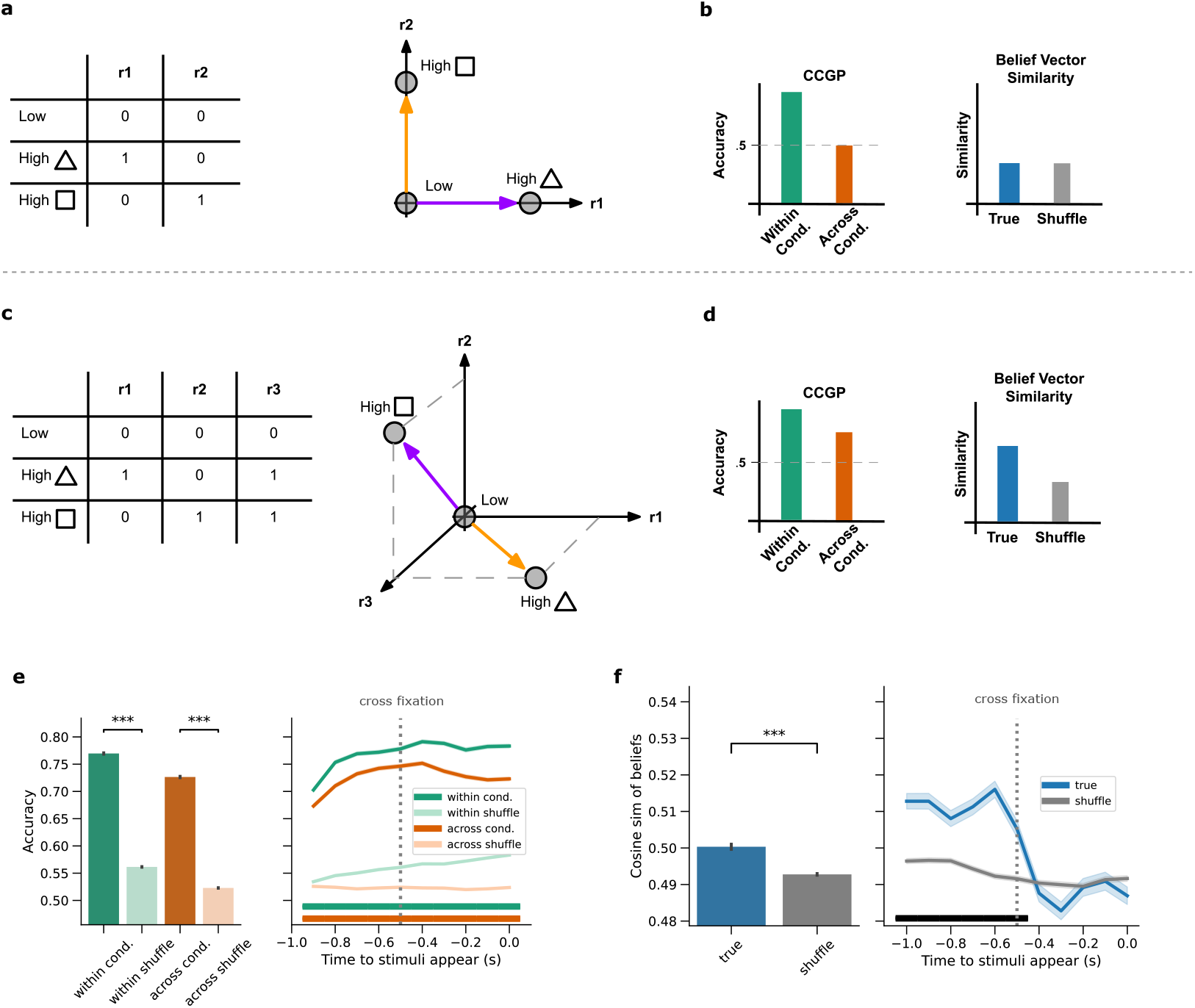
Belief representations are aligned in neural activity space. **a-d**, Alternative hypotheses for the geometric organization of belief representations. **a**, Toy example illustrating orthogonal belief encoding across two features (square and triangle) using two neurons. Feature-specific firing rates (left) produce orthogonal dimensions in neural activity space (right). **b**, Predicted outcomes under the orthogonal hypothesis for cross-condition generalization performance (CCGP, left) and cosine similarity analysis (right). Orthogonal belief representations yield poor cross-condition accuracy and near chance belief vector similarity. **c**, Same as **a**, for aligned belief representations. A third, confidence-selective neuron (encoding confidence independent of feature identity) produces partial alignment along one dimension. **d**, Predicted outcomes under the aligned hypothesis. Aligned representation yield above-chance cross-condition accuracy (left) and above-chance belief vector similarity (right). **e, f**, CCGP and similarity analysis applied to recorded population neural activity during the 1s pre-stimulus interval, aggregated across the interval (left) and split by time bin (right). Stars (left panels) and horizontal bars (right panels) represent significance levels relative to corresponding shuffles: *** and thick: *p* < 0.001, ** and moderate: *p* < 0.01, * and thin: *p* < 0.05

To probe this question, we focused on pairs of features *X* and *Y*, and partitioned the model-predicted belief states into three categories: low confidence (Low), high confidence preferring feature *X*, (High *X*), and and high confidence preferring feature *Y* (High *Y*). Because different prior beliefs may drive distinct search strategies and stimulus choices once stimuli appear, we focused on the 1-second pre-stimulus interval, as in previous analyses.

Both organizational schemes preserve the linear separability of confidence and preference as previously defined, so decodability of the two variables alone would not reveal the organization. To assess the alignment of belief representations, we applied a variant of cross-condition generalization performance (CCGP) analysis [29–31], (Methods 5.12). Under the orthogonal hypothesis, a decoder trained to discriminate belief states for one feature (ex. Low vs. High square) would perform well within those conditions but would not generalize to another feature (ex. Low vs. High triangle) (Fig 8b, left). Under the aligned hypothesis, however, such a decoder would generalize across features (Fig 8d, left). Applying CCGP to the recorded neural population revealed high cross-condition decoding accuracy that is significantly above chance across all time bins (Fig 8e), indicating aligned belief representations. To complement this analysis, we examined the cosine similarity between the averaged population activity vectors representing beliefs about different features (Figs 8a,c purple and orange arrows). Under the orthogonal hypothesis, true cosine similarity between population vectors for different feature beliefs would not differ from a shuffled baseline (Fig 8b, right). However, under the aligned hypothesis, true cosine similarity would significantly exceed the shuffled baseline (Fig 8 d, right). In our data, true cosine similarity between belief vectors consistently exceeded the shuffled baseline across most time bins (Fig 8f), providing further evidence for beliefs being aligned in neural activity space. This aligned representation was present across all recorded brain regions (Fig S7).

### 2.6 Belief representations are organized by feature dimensions

In our task, features are drawn from three feature dimensions: shape, color and texture. Each stimulus can only contain one feature from each category. Does this feature structure and statistical constraint influence the organization of feature representation? To 18 study this, we similarly examine pairs of features at a time and their corresponding belief partitions, as in the previous analysis. We can additionally group pairs of features as being either within-dimension (e.g. square vs. triangle) or cross-dimension (e.g. square vs. green).

If monkeys’ strategy involves focusing attention on a single dimension at a time and then deciding which of the three options to choose [11, 32, 33], one might expect that beliefs for within-dimension feature pairs are more aligned than across-dimension feature pairs (Fig 9a). If this were the case, cross-condition accuracy from CCGP would be higher for within-dimension pairs than across-dimension pairs, and the cosine similarities of belief vectors between within-dimension pairs would be higher than those between cross-dimension pairs (Fig 9b).

**Fig. 9:**
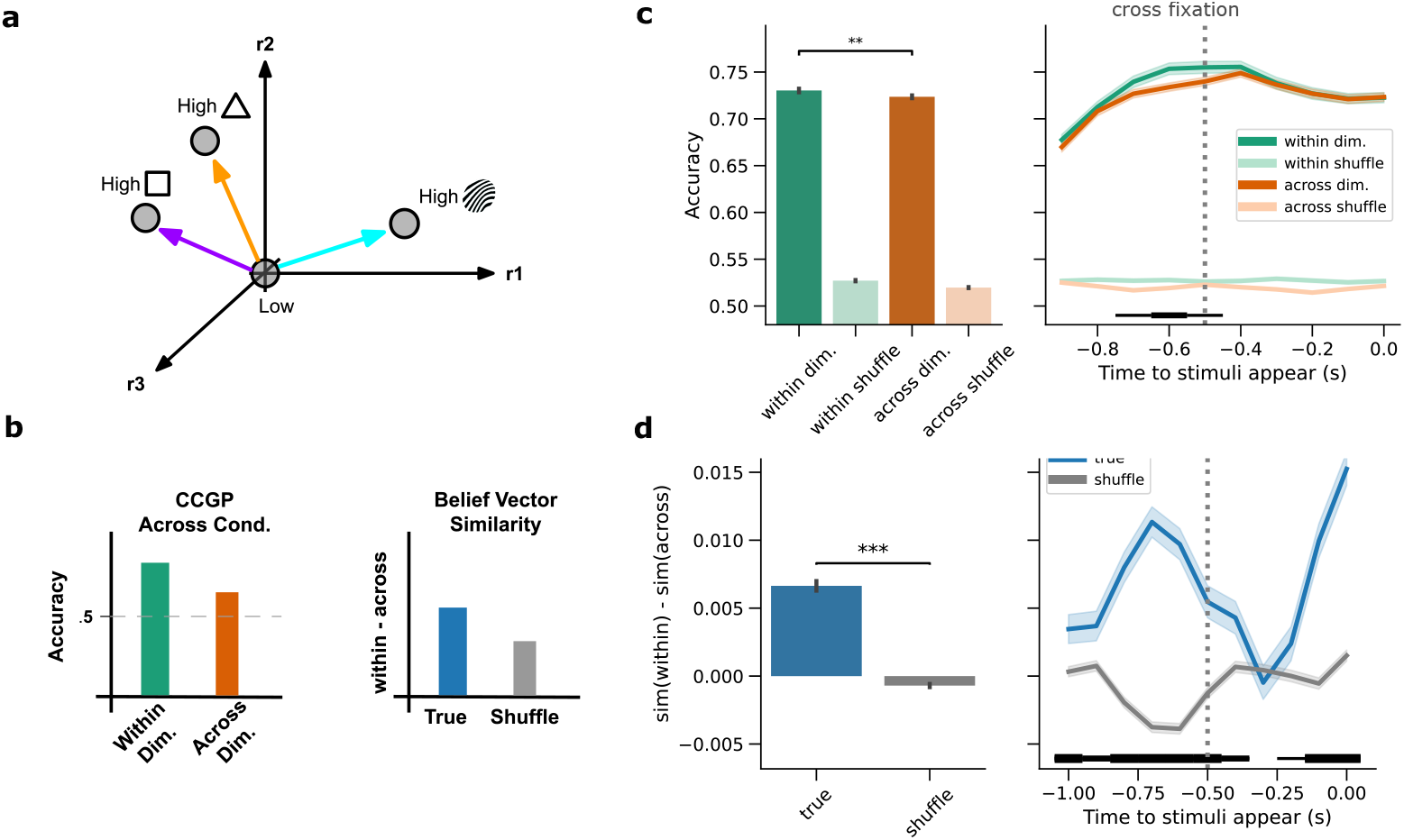
Beliefs are organized by feature dimensions **a**, A toy example depicting a within-dimension organization, where beliefs of features in the same dimension (e.g. square vs. triangle) are more aligned than beliefs of features in different dimensions (e.g. square vs. swirl). **b**, Predicted outcomes for cross condition generalization performance (CCGP, left) and similarity analysis (right). Within-dimension organization will lead to higher across-condition accuracy for pairs of within dimension features than across dimension features, and a corresponding difference in belief similarities between within and across dimension pairs to be higher than chance. **c, d**, Within and across dimension CCGP and similarity analysis applied to recorded population neural activity during the 1s pre-stimulus interval, aggregated across the interval (left) and split by time (right). Stars (left panels) and horizontal bars (right panels) represent significance levels relative to corresponding shuffles: *** and thick: *p* < 0.001, ** and moderate: *p* < 0.01, * and thin: *p* < 0.05

In the recorded population, we find that cross-condition accuracy from CCGP is indeed slightly but significantly higher for within-dimension feature pairs than cross (Fig 9c), while the difference in cosine similarities of belief vectors between within-dimension pairs and across-dimension pairs is significantly higher than a shuffle (Fig 9d), providing evidence towards a within-dimension belief organization in the brain. A within-dimension organization is present across most regions of interest (Fig S8).

## 3 Discussion

Our work uncovers broadly distributed neural representations of beliefs and other task variables across multiple brain regions. Trial-by-trial, belief-related neural activity evolves in a manner consistent with Bayesian inference, with updates influenced not only by trial outcomes but also by prior belief states. We further quantify the geometry of belief representations and find an organization that is non-orthogonal and structured according to visual feature dimensions.

### 3.1 Distributed implementation of belief-based inference

We studied belief representation using a multi-dimensional inference task, where presented stimuli are high-dimensional but can be decomposed into a low-dimensional feature set that governs reward. In such a task, traditional action-value based RL algorithms used in past studies [34–37] would suffer from the curse of dimensionality [38], as tracking values across individual stimuli leads to poor performance (Fig 1f). Instead, our behavioral modeling suggests that monkeys leverage an internal model of the task to decompose stimuli into visual features and track values or beliefs across these features.

Previous work has proposed various formulations for such internal models. Feature-based decomposition has been proposed to arise through attentional mechanisms, [11, 32, 39, 40], where top-down processes selectively guide attention toward certain features of a visual scene. Others describe the internal model as a “cognitive map” that captures transitions between abstract task states [41–44]. In the Bayesian formulation that guides our neural analyses [24, 25, 45], the internal model is a generative model of the task that is used when updating beliefs about hidden states. While past work has quantified which form of internal model best describes behavior in similar tasks [39, 46–48], more careful analysis of our specific paradigm would be required to strictly characterize the monkeys’ internal model and distinguish among these alternatives. Nonetheless, feature RL models and Bayesian belief state models make quite similar predictions for hidden internal variables (Figs S1b,f), and share the core idea of a decomposed feature representation.

We find that representations of task variables and beliefs are broadly distributed across regions, aligning with recent findings of brain-wide, distributed coding of task-relevant cognitive variables [49–52]. This distributed representation may highlight the diversity of roles that beliefs play in task performance: Beliefs guide decision-making, visual search strategy, and feedback integration. The region-specificity of when confidence and preference signals peak and remain stable may illuminate the distinct computational contributions of each region to solving the task.

### 3.2 Geometry of belief representations

Our analysis of representational geometry uncovered an aligned, non-orthogonal structure in which beliefs about different features share common representational dimensions. This alignment may reflect shared neural mechanisms for tracking belief confidence, independent of specific feature identity. Similar “abstract” representational geometries have been reported in other cognitive tasks [29–31, 53]. One possible explanation for this organization is that increases in confidence about the hidden rule, regardless of the underlying feature, may trigger common neural processes such as heightened attention and task engagement, or shifts from exploratory to exploitative search strategies during decision-making [54, 55].

We further find that belief representations are organized by visual feature dimension (color, shape, pattern). This organization could arise through two distinct mechanisms. First, belief representation may inherit the semantic organization as visual feature representations, where different visual feature dimensions are encoded independently in visual cortex [56, 57]. Alternatively, the feature-dimension organization of beliefs may reflect a hierarchical inference strategy in which monkeys first track beliefs across feature dimensions to guide visual attention during search, then make decisions about specific features within the attended dimension. [11, 32, 33]. Distinguishing between these hypotheses would require additional task designs that decouple visual feature structure from the inference hierarchy, along with more detailed behavioral analysis of search patterns and reaction times.

## 4 Extended Figures

**Fig. S1:**
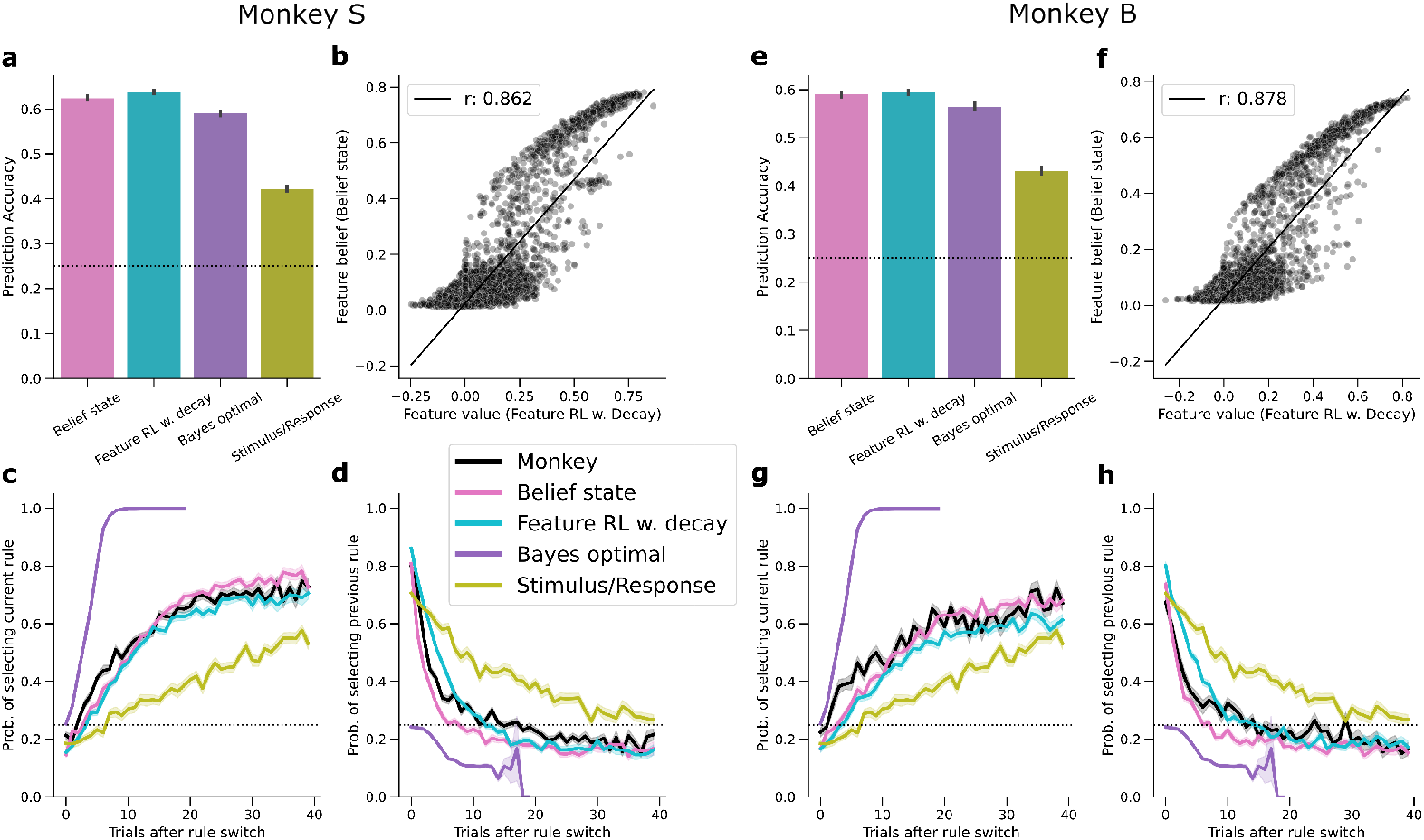
Comparison of behavioral models to monkey data. **a, e**, Prediction accuracies of monkeys’ stimuli selections for each model, evaluated on held-out test sessions for Monkeys S, B, respectively. **b, f**, Correlation between inferred feature values (from the feature RL w. decay model) and feature beliefs (from the belief state model). Scattered circles indicate feature/trial pairs, randomly sub-sampled to 10,000 pairs for display. Line indicates best-fit linear regression line, reporting Pearson’s r coefficient in the figure legend. **c, g**, Probability of selecting stimuli containing the current rule after rule switches, for both monkeys S and B in black, and different fitted models. Fitted models were run on 50 sessions of 1000 trials each. Shaded regions indicate s.e.m. **d, h**, same as **c, g** but for probability of selecting a stimulus containing the previous rule.

**Fig. S2:**
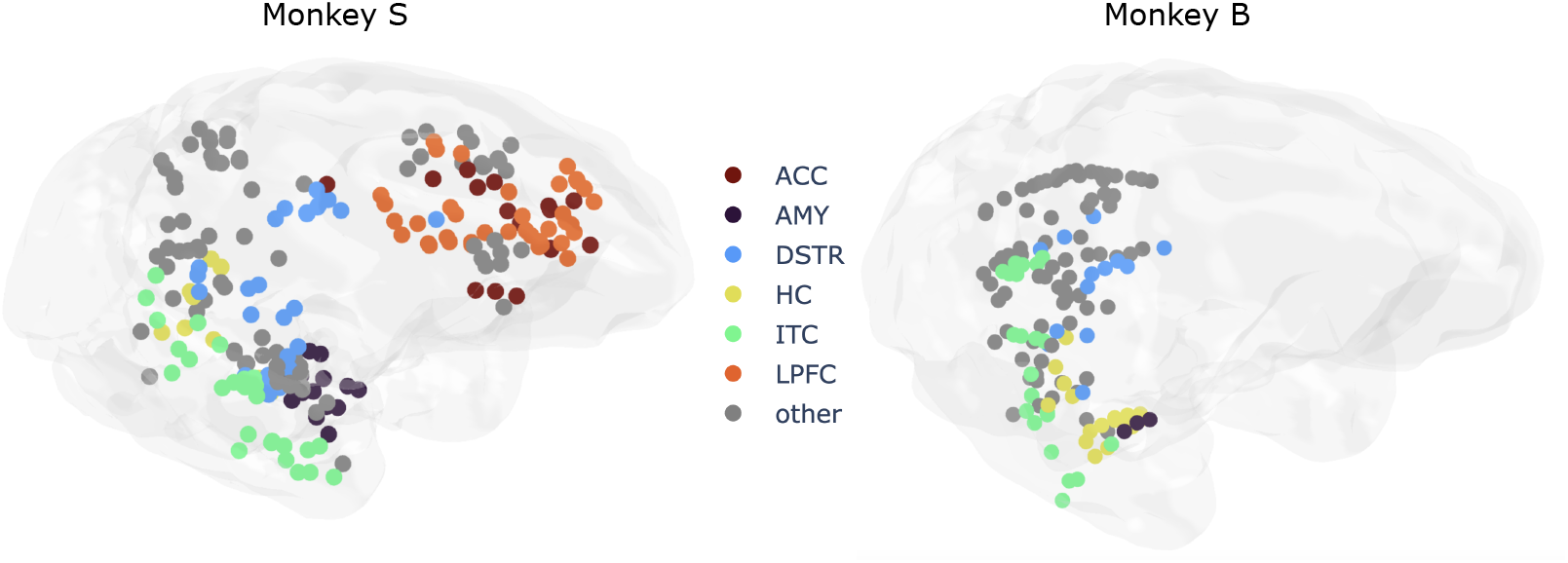
Recorded single unit positions for Monkey S (left) and Monkey B (right). Colors indicate recorded regions, with grey indicating units that did not fall into an individually analyzed region.

**Fig. S3:**
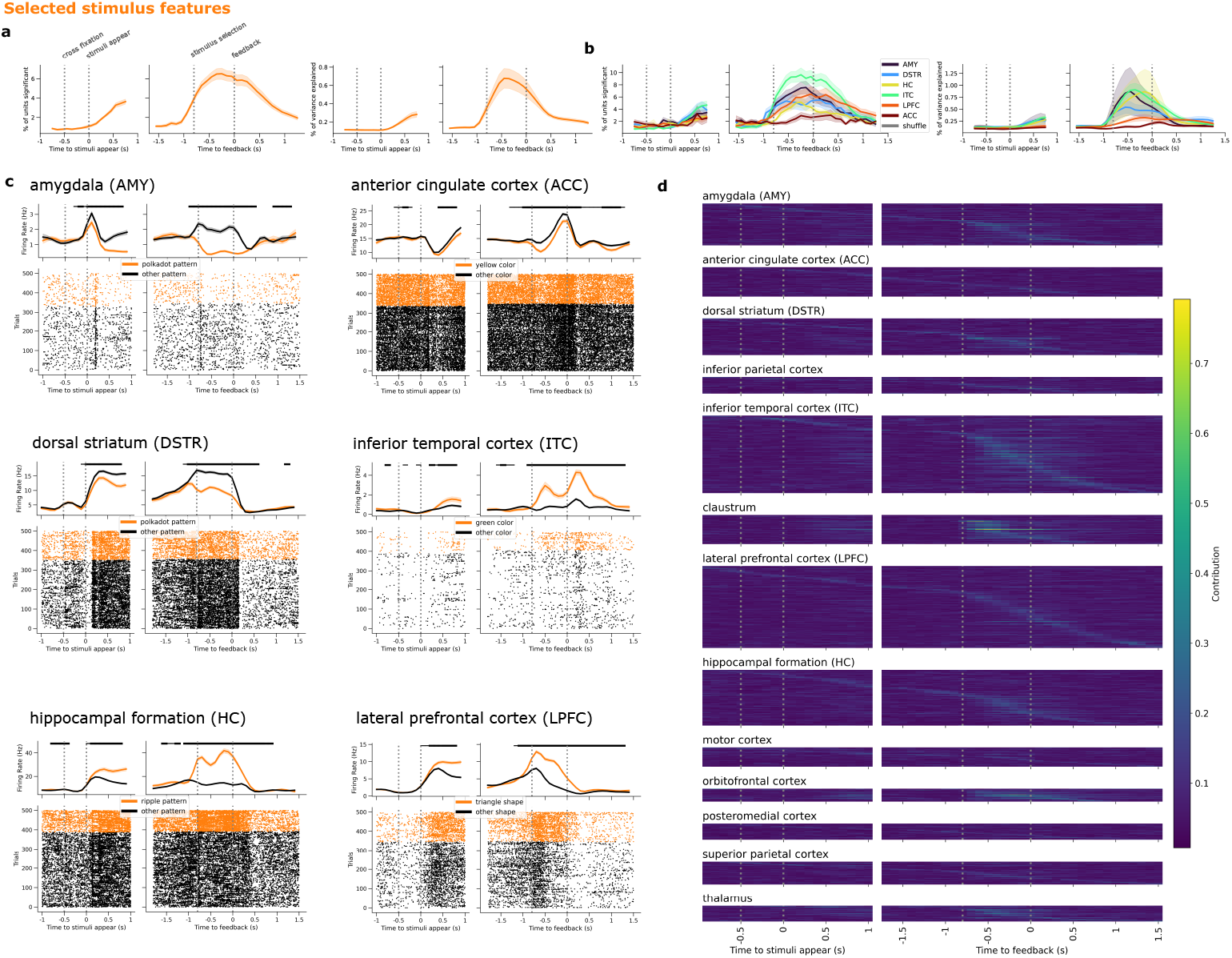
Additional analyses of selected stimulus feature representations. **a** left, percentage of units in the entire recorded population significantly encoding selected stimulus features across time, assessed via ANVOA with a 500ms sliding window of activity, *p* < 0.01. right, percentage of variance in firing rates explained by selected stimulus features d. **b**, same as a, but split by regions of interest. **c** PSTHs and spike rasters of example selected stimulus feature selective neurons in each region. Horizontal black bar thickness indicates significance level: thick: *p* < 0.01, thin: *p* < 0.05. **d**, Single unit contributions to the whole population selected stimulus feature decoders, split by regions. Brighter cells indicate a larger decoder weight magnitude. Per-region, rows are sorted by the time bin of peak contribution.

**Fig. S4:**
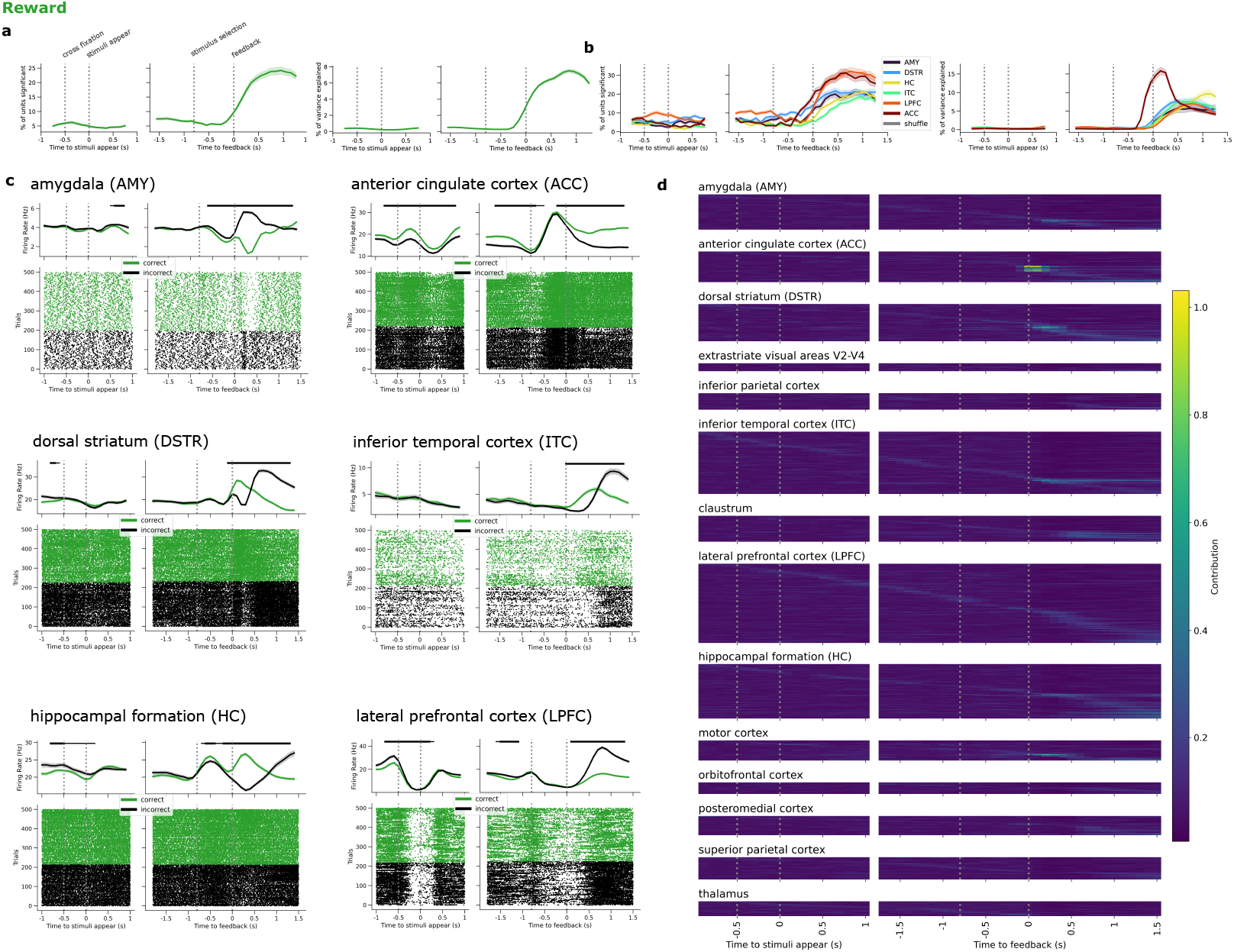
Additional analyses of reward representations. **a** left, percentage of units in the entire recorded population significantly encoding reward across time, assessed via ANVOA with a 500ms sliding window of activity, *p* < 0.01. right, percentage of variance in firing rates explained by reward d. **b**, same as a, but split by regions of interest. **c** PSTHs and spike rasters of example reward selective neurons in each region. Horizontal black bar thickness indicates significance level: thick: *p* < 0.01, thin: *p* < 0.05. **d**, Single unit contributions to the whole population reward decoders, split by regions. Brighter cells indicate a larger decoder weight magnitude. Per-region, rows are sorted by the time bin of peak contribution.

**Fig. S5:**
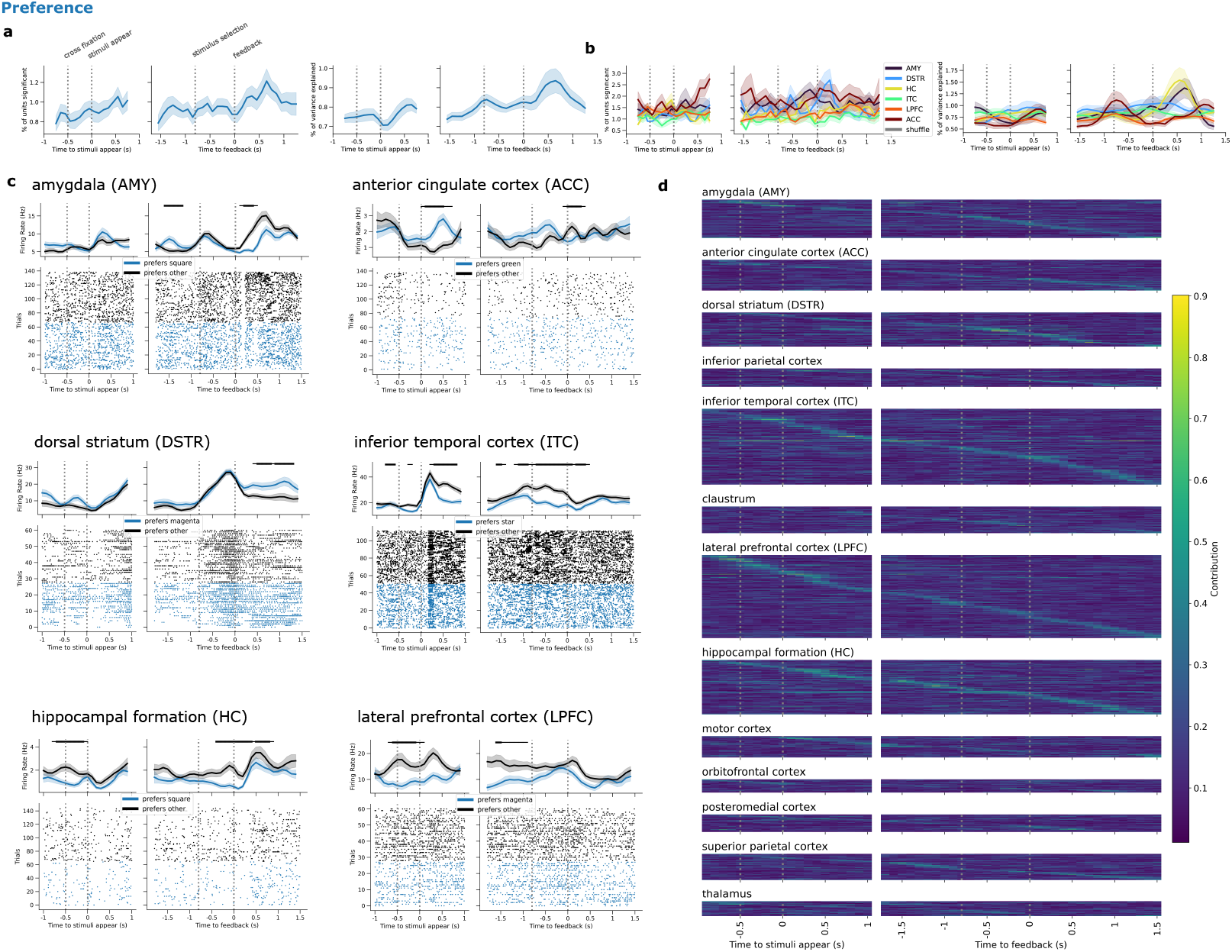
Additional analyses of preference representations. **a** left, percentage of units in the entire recorded population significantly encoding preference across time, assessed via ANVOA with a 500ms sliding window of activity, *p* < 0.01. right, percentage of variance in firing rates explained by preference d. **b**, same as a, but split by regions of interest. **c** PSTHs and spike rasters of example preference selective neurons in each region. Horizontal black bar thickness indicates significance level: thick: *p* < 0.01, thin: *p* < 0.05. **d**, Single unit contributions to the whole population preference decoders, split by regions. Brighter cells indicate a larger decoder weight magnitude. Per-region, rows are sorted by the time bin of peak contribution.

**Fig. S6:**
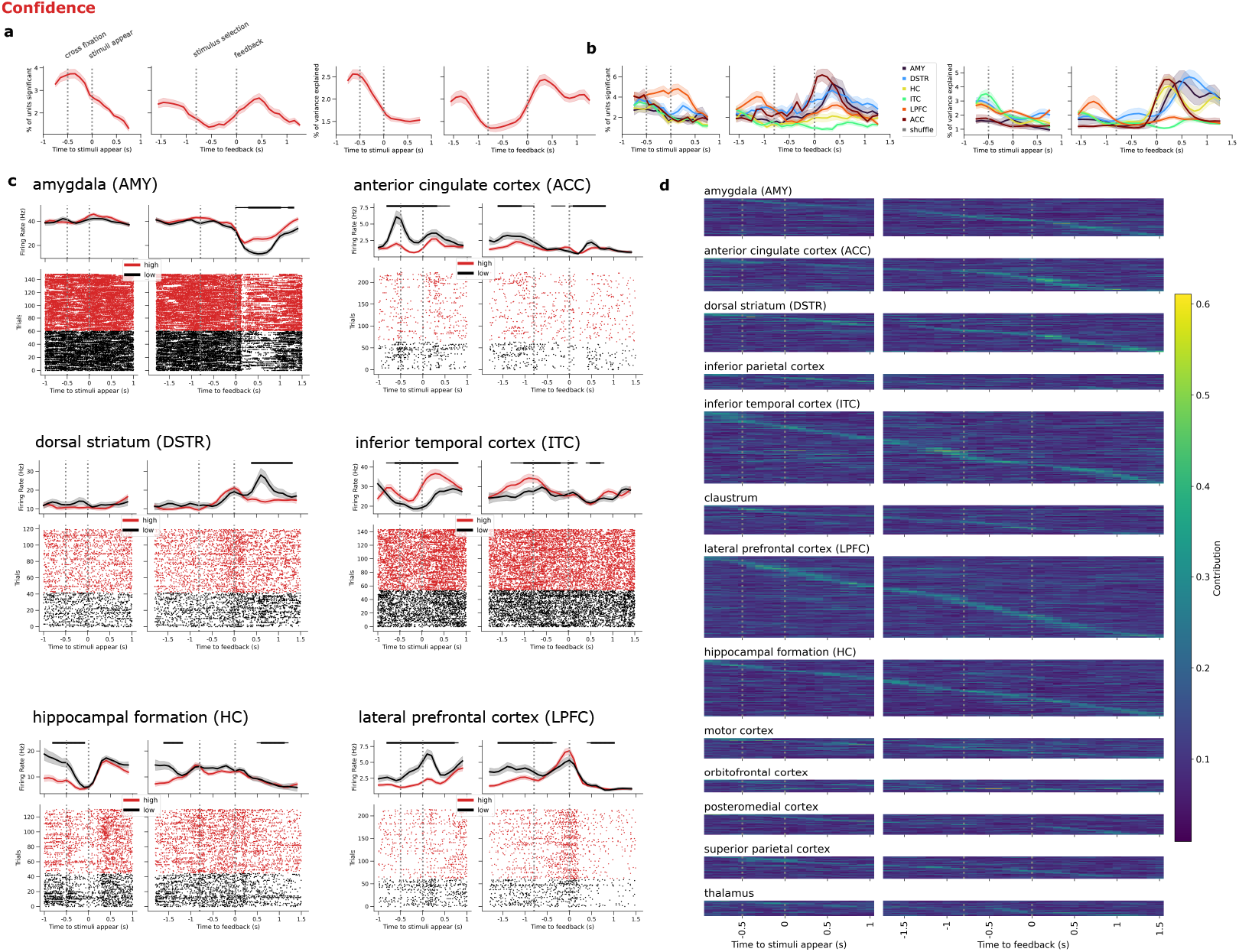
Additional analyses of confidence representations. **a** left, percentage of units in the entire recorded population significantly encoding confidence across time, assessed via ANVOA with a 500ms sliding window of activity, *p* < 0.01. right, percentage of variance in firing rates explained by confidence d. **b**, same as a, but split by regions of interest. **c** PSTHs and spike rasters of example confidence selective neurons in each region. Horizontal black bar thickness indicates significance level: thick: *p* < 0.01, thin: *p* < 0.05. **d**, Single unit contributions to the whole population confidence decoders, split by regions. Brighter cells indicate a larger decoder weight magnitude. Per-region, rows are sorted by the time bin of peak contribution.

**Fig. S7:**
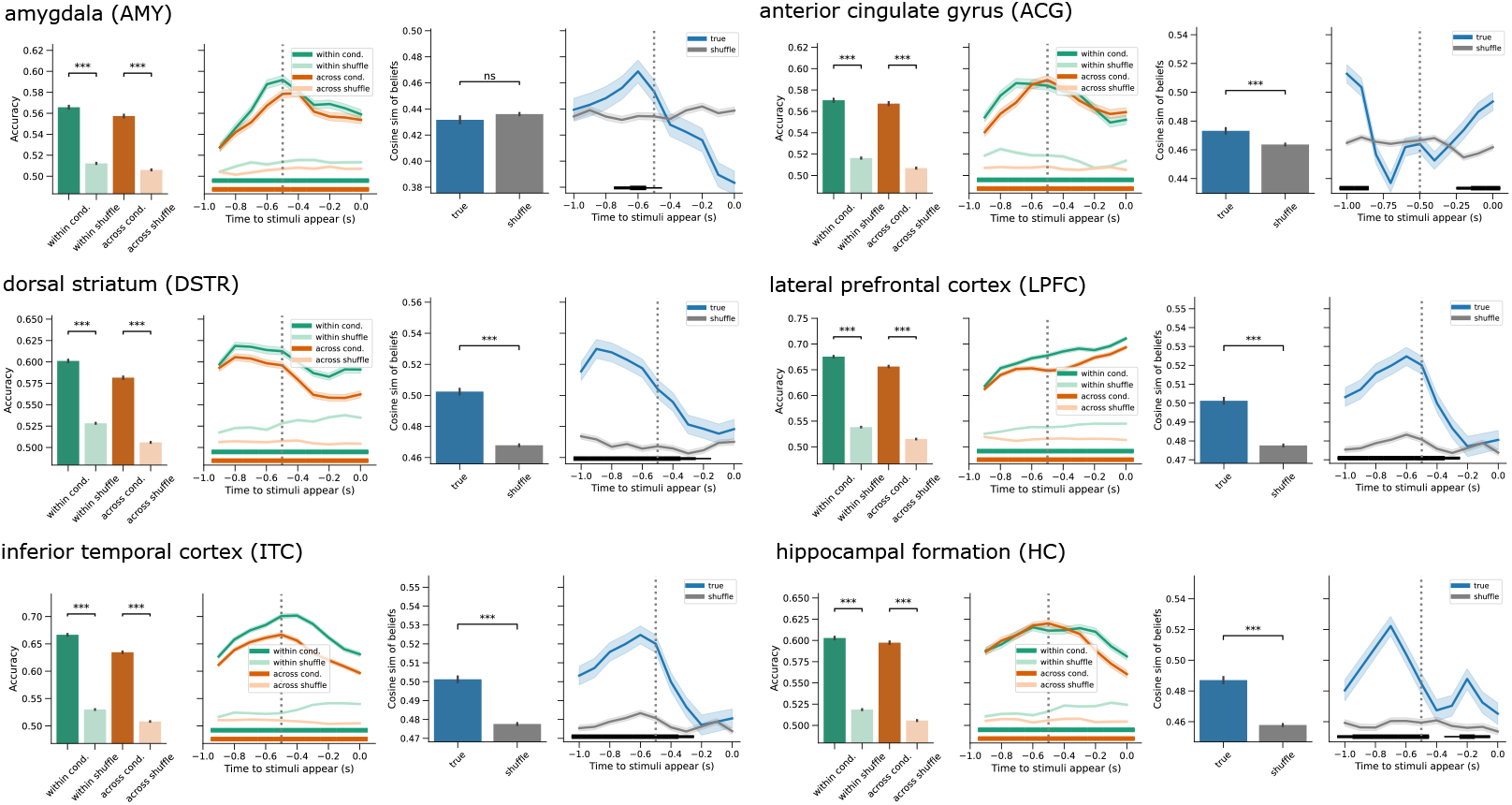
CCGP and similarity analysis applied per-region to recorded population neural activity 1s before stimuli appearing, aggregated across the interval (bar plots) and split by time (line plots). Stars in the bar plots and horizontal bars in the line plots represent significance levels relative to corresponding shuffles: *** and thick: *p* < 0.001, ** and moderate: *p* < 0.01, * and thin: *p* < 0.05

**Fig. S8:**
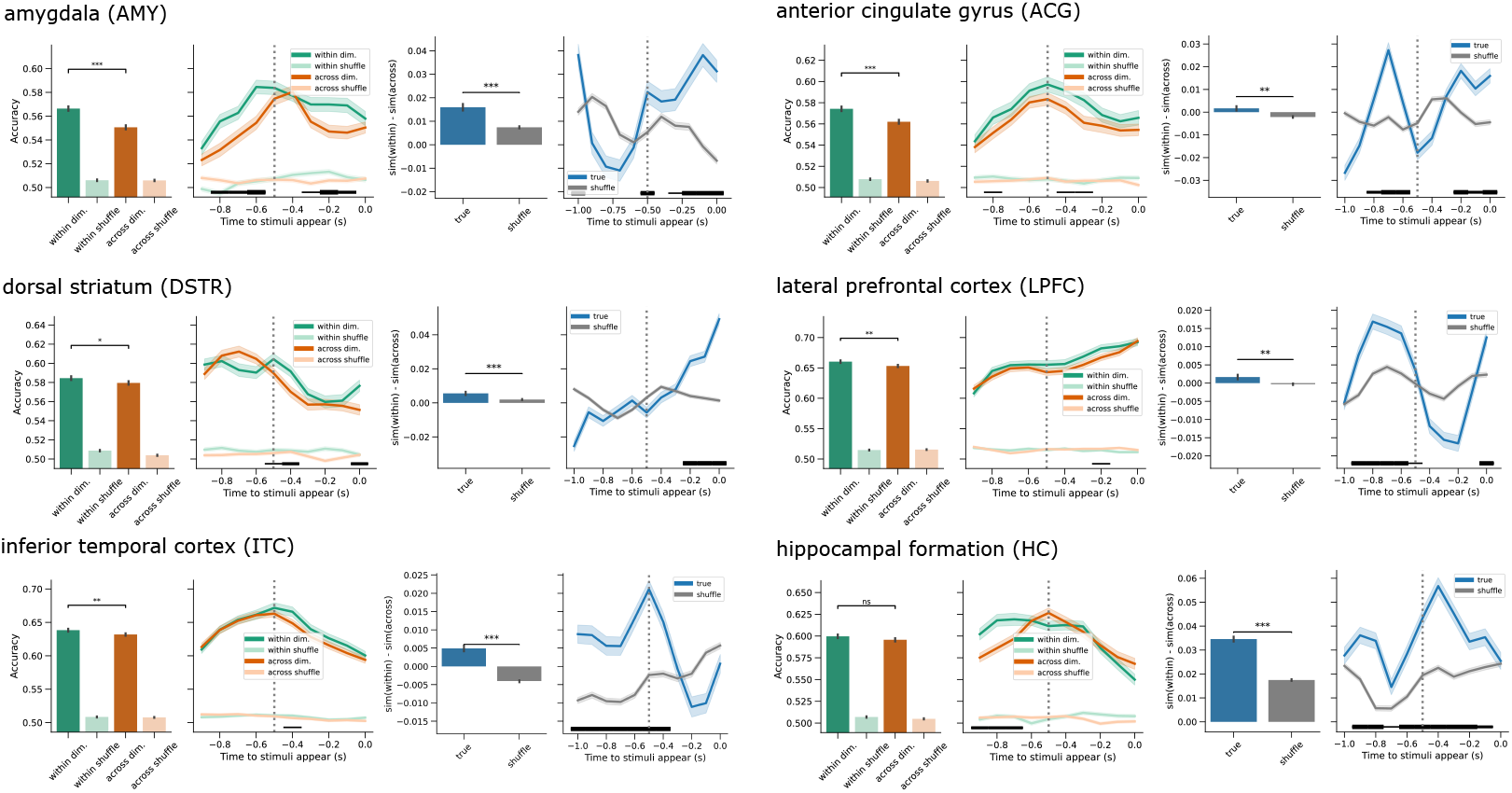
Within and across dimension CCGP and similarity analysis applied per-region to recorded population neural activity 1s before stimuli appearing, aggregated across the interval (bar plots) and split by time (line plots). Stars in the bar plots and horizontal bars in the line plots represent significance levels relative to corresponding shuffles: *** and thick: *p* < 0.001, ** and moderate: *p* < 0.01, * and thin: *p* < 0.05

## 5 Methods

### 5.1 Monkeys

All procedures were carried out in accordance with the National Institutes of Health guidelines and were approved by the University of Washington Institutional Animal Care and Use Committee. Electrophysiological recordings were conducted in two adult female rhesus monkeys (Macaca mulatta, referred to as “monkey B” and “monkey S”). The majority of monkeys’ caloric intake during testing was provided in the form of the food slurry reward. Monkeys were supplemented with fruits and vegetables as well as monkey chow after testing each day. Daily pretesting weights were taken to monitor weight, and the caloric intake was adjusted to maintain a vet-approved weight range based on sex and age. Both animals had ad libitum access to drinking water.

### 5.2 Task Description

Following the calibration task, monkeys performed the multi-dimensional inference task. The monkey initiated each trial by visually fixating on a white cross (0.5°) at the center of a gray screen. After 500 ms of successful fixation, the cross disappeared and was replaced by an array of four stimuli, which stayed on the screen during the following self-paced decision period. The monkey was free to explore the array for up to 4 s; she signaled her choice of a stimulus by maintaining her gaze within a 9×9° window centered on the stimulus for 800ms. Immediately following a successful choice, the monkey received feedback. If the stimulus containing the current rule feature was selected, the monkey received a food slurry reward over 1.4 s; if the chosen stimulus did not contain the target feature, or if she failed to make a choice within 4 s, a timeout period (either 1 or 5 s) occurred, consisting of the presentation of a blank gray screen and no reward. (The 1s timeout period was used for Monkey B to increase the number of trials completed per session, after confirmation that this reduction did not alter her trial-to-criterion performance.) A 400 ms intertrial interval (ITI) immediately followed the feedback period. Rule switches were based on performance: Once the monkey made 8 consecutive correct responses or 16 correct responses in 20 or fewer trials, the current rule feature was changed to a new rule feature. See [23] for a detailed description of the training steps used to shape monkeys’ behavior on this task. During testing, monkeys were seated and head-fixed 60 cm from a 19-inch 800×600 pixel CRT monitor displaying images at a refresh rate of 120 Hz noninterlaced. At this distance, the monitor screen subtended 33 by 25 degrees of visual angle. Eye movements were recorded using a noninvasive infrared eye-tracking system (EyeLink 1000 Plus, SR Research). The task was presented using experimental control software (NIMH Cortex). Calibration of the infrared eye-tracking system was accomplished using a multi-point manual calibration task designed to accurately map eye movements to positions across the entire screen.

### 5.3 Electrophysiological Recordings

Before implantation of recording hardware, we collected T1-weighted MRI scans of each monkey to localize brain regions of interest and to guide placement of custom-built recording chambers. Prior to the placement of recording hardware, a titanium post for holding the head was implanted. Recording hardware consisted of large-scale semi-chronic microdrive recording systems developed in the lab of Charles Gray [58]. Both monkeys were implanted with microdrive systems containing 124 independently moveable tungsten microelectrodes (FHC Inc., Bowdoinham, Maine, USA; Alpha Omega Co. USA Inc., Norcross, Georgia, USA) positioned over the temporal lobe, centered on the hippocampal formation. Monkey S had an additional microdrive system containing 96 microelectrodes positioned over the frontal lobe. All microdrives remained on the animals for the duration of the experiment. After chamber placement, a second MRI scan was performed with MRI-visible fiducial markers mounted on each animal’s head. We generated .stl models using a combination of pre- and post-operative MRI scans and 3D scans of the animals’ heads; we then used 3D Slicer (https://www.slicer.org) to align these models to estimate the placement and angle of the microdrives and the electrode starting positions.

After drive placement and throughout the experimental timeline, electrodes were advanced through precise manual turns of the leadscrew attached to the actuator mechanism of each microwire; the distance correlation of a full turn of the screw was measured during non-implanted practice turns. We initially advanced all electrodes a minimal distance into the brain during surgery, then each electrode was incrementally advanced over the course of recordings. For each recording session, a subset of electrodes was manually advanced to desired recording locations. We kept a careful log of the turn counts for each electrode, allowing us to estimate the total electrode distance traversed across recording sessions. These distances were transformed into 3D coordinates of recording locations using custom MATLAB code, with the aligned .stl models used for reference. To assign these coordinates to brain regions, we processed the T1-weighted MRI scans using methods described in Jung et al. to automatically segment the brain. We used AFNI’s @animal_warper to perform skull-stripping and to align the brain to the macaque NMT v2 Atlas, then performed parcellation to apply labels to each brain region according to the cortical hierarchy atlas of the rhesus monkey (CHARM) [59] and the subcortical hierarchy atlas of the rhesus monkey (SARM) [60]. Neural data acquisition was performed using a Neuralynx Cheetah system (version 6.3.1). Neural signals from each wire electrode were captured at 32 kHz and collected in wideband format (0.1 – 8000 Hz). Spike components were extracted from the wide-band extracellular signal using post hoc processing in MATLAB. The wideband signal was high-pass filtered using a fourth-order Butterworth filter with a cutoff frequency of 400 Hz, applied in zero-phase (`filtfilt`) mode to prevent phase distortion. Candidate spikes were captured by identifying threshold crossings of the filtered signal. Non-biological waveform shapes and simultaneous spikes across channels were eliminated via an automated heuristic method prior to sorting. The branch cluster method from ISO-SPLIT 5 [61] was used to identify and separate spike waveform clusters. Further manual curation was used to verify accurate identification of spike vs. noise clusters. In all subsequent analyses, we examine neural activity around two trial intervals, one 1000ms before/after stimulus onset, and one 1800ms before feedback onset, to 1500ms after. Across both intervals, we computed per time-bin firing rates by binning spikes into 100ms bins, and smoothing with a Gaussian kernel of 100ms. We further identified and excluded neurons exhibiting slow drifts in firing rate across the session. For each neuron, per trial, we first computed average firing rates across both trial intervals. We then computed the power spectral density (PSD) of these trial-averaged firing rates across trials. Neurons were excluded if more than 10% of the total spectral power was concentrated in low-frequency components, corresponding to periods longer than 100 trials, indicating slow temporal drift in firing rate.

### 5.4 Behavioral models

To explain monkey behavior on the task, we compared and tested several behavioral models. To describe the models in detail, we start with framing the task in consistent notation.

In our task, stimuli are composed of color, shape and pattern features, which we describe via three feature sets:

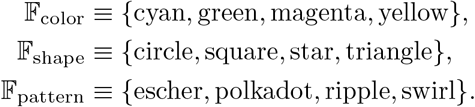

On each trial *k*, the hidden rule feature *z*_*k*_ may be any color, shape or pattern, such that:

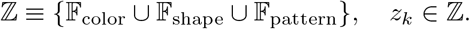

Per-trial *k*, the monkey sees four stimuli on a screen, each of which has a color, shape and pattern, with the *i*-th stimulus denoted as:

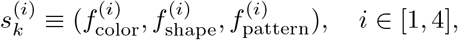

with:

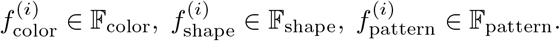

There are no feature repeats across stimuli, such that:

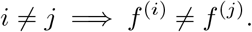

Additionally, all twelve features are all present on the screen, such that:

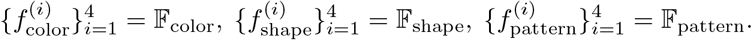

On each trial the monkey makes a selection 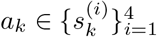 and correspondingly receives reward *r*_*k*_ ∈ {0, 1} for incorrect/correct feedback respectively, with reward governed by whether the selected stimulus contains the hidden rule feature:

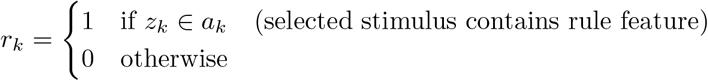

Any behavioral model should provide per-trial probabilities of selecting each presented stimuli:

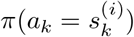

#### 5.4.1 Bayes Optimal model

We start with formulating how an monkey should optimally perform the task. We assume that on every trial, the monkey has learned to track a belief state *b*_*k*_(*z*), a posterior probability distribution of the hidden rule feature *z* given a history of past stimuli selections, and rewards: *b*_*k*_(*z*) ≡ *p*(*z* |*h*_*k*_), where history is defined the sequence of past stimuli selections and rewards:

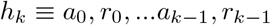

The task is Markovian in nature, so the monkey’s belief state may be recursively updated trial-by-trial via Bayesian inference, such that the belief of rule *z* on trial *k* + 1 is computed via:

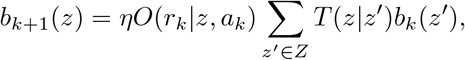

where *η* is a normalization constant, *O*(*r*_*k*_|*z, a*_*k*_) is a reward observation probability, *T* (*z* |*z*^*′*^) is a rule transition probability from a previous rule *z*^*′*^ to current rule *z*, and *b*_*k*_(*z*^*′*^) is the prior belief of *z*^*′*^ on trial *k*. For an optimal performer, the reward observation and rule transition probabilities will match the task parameters exactly, such that:

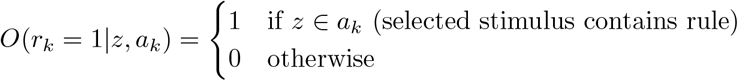

As described in 5.2, rule transitions depend on the monkey’s performance. For notational simplicity, we will let *p*(*k*) = 1 to denote a trial *k* where the rule switching criteria has been satisfied, and *p*(*k*) = 0 for trials otherwise. The full rule transition probability can then be expressed as:

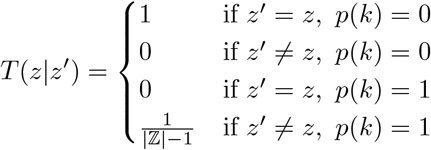

where |ℤ|= 12. With these feature beliefs, a monkey can than compute the probability that each stimuli contains the rule on each trial:

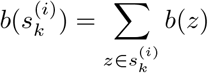

The monkey then can select the stimulus with the highest probability of containing the rule:

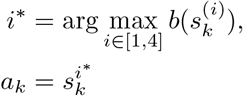

Since this model was used to describe optimal behavior, it contains no free parameters and was not directly fit to monkey behaviors.

#### 5.4.2 Belief state model

To account for the fact that monkey performances are suboptimal, we consider the possibility that the monkey’s internal model of the task differs from the true task specifications [25]. As in 5.4.1, we still assume that monkeys are tracking beliefs in rules per-trial, *b*_*k*_(*z*), and updating them via Bayesian inference. However, we choose a specific parametrization of the reward observation probability *O* and rule transition probability *T*, such that they may differ from the true task. Specifically, we consider reward observation probabilities of the form:

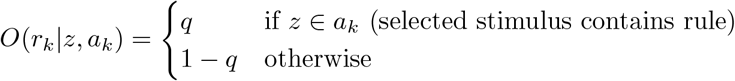

where *q* ∈ [0, 1] is a free parameter. Though the true rule transition in the task is performance-based, here we make the simplifying assumption of a stationary transition probability of the form:

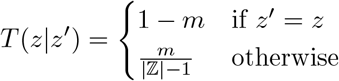

where *m* ∈ [0, 1] is a free parameter as well. Additionally, we assume that given a belief *b*_*k*_(*z*) and a set of stimuli on the screen, the monkeys are performing a selection strategy closer to probability matching, where the probability of selecting any particular stimulus on the screen is given by the policy *π*(*a*_*k*_|*b*_*k*_), computed as:

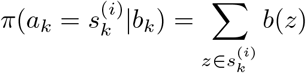

We initialize belief states with uniform probability across all possible features:

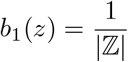

Altogether, the belief state behavioral model is a two parameter model, with free parameters *θ* = {*q, m*}.

#### 5.4.3 Feature RL with Decay

Following past work modeling a similar task [39], we additionally compare a behavioral model which draws inspiration from Q-learning in reinforcement learning [38]. In this model, on each trial *k*, monkeys are assumed to track a scalar value for each possible rule feature *z* denoted here as:

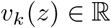

During stimulus selection, monkeys integrate feature values into stimulus values 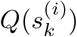, based off the features present on each stimulus:

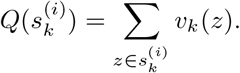

Per-trial stimulus selections *a*_*k*_ are then made probabilistically in proportion to the stimulus values, using a softmax function with temperature parameter *β* ∈ (0, ∞):

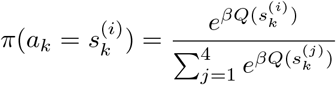

After feedback, monkeys update features of the selected stimulus via a reward prediction error (RPE) computed from the selected stimulus value and learning rate *α* ∈ [0, 1]. Additionally, to account for finite memory, the values of features not present on the selected stimulus are also decayed with a rate *d* ∈ [0, 1]. Altogether, feature values are updated per-trial via:

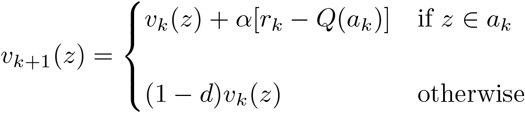

We initialize feature values as 0 across all features:

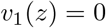

The Feature RL with Decay model is a three parameter model, with free parameters *θ* = {*β, α, d*}.

#### 5.4.4 Stimulus/Response Mapping

One possibility is that monkeys may have performed the task by simply tracking which stimuli led to reward, ignoring the task’s feature-based structure. To investigate this, we can assume that monkeys are perfectly tracking binary values of whether a reward was last received when selecting a particular stimulus. Across all the 64 possible stimuli, this can be denoted as:

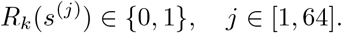

After feedback on each trial, only the selected stimulus gets updated:

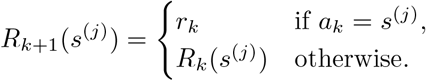

During selection, previously rewarded stimuli are selected preferentially with:

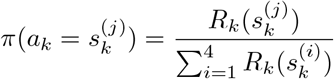

With uniform probability selection if none of the presented stimuli were previously rewarded. Like the Bayes Optimal model, this model also contains no free parameters and was not directly fit to monkey behaviors.

#### 5.4.5 Model Fitting

To fit both the Belief State and Feature RL with Decay models to monkey behavior, we find parameters *θ*^∗^ which maximize the likelihood of observing the monkey stimulus selections *â*_*k*_ from the model on each trial:

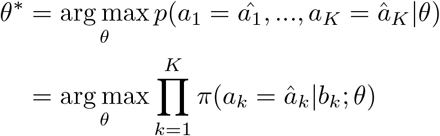

Models were fitted per monkey, using the `scipy.optimize.minimize` function in Python, with the Nelder-Mead optimization method. After uncovering best-fit model parameters per-monkey, per-trial beliefs states for the Belief State model were then recursively inferred using the monkey’s per-trial stimuli selection *â*_*k*_ and reward 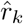:

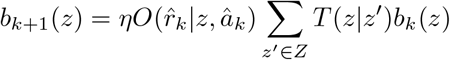

Feature values for the Feature RL with Decay model were similarly obtained:

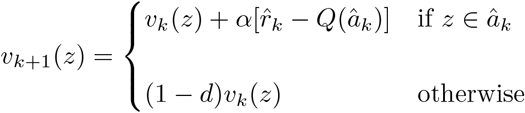

### 5.5 Partitioning Belief States, Session Selection

The per-trial belief states inferred from our behavioral model correspond to points on a twelve-dimensional probability simplex, requiring a high number of samples to sufficiently study the neural representation of this space. To address this difficulty, we partition our belief states into discrete partitions, and examine beliefs either a feature at a time, or pairs of features at a time.

For a given belief state distribution *b*(*z*), we can first define a preferred feature *z*^∗^ as the feature with the current highest belief:

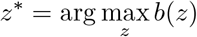

If analyzing beliefs a feature at a time, relative to some feature *X* of interest, we can then partition all possible belief states into three partitions:

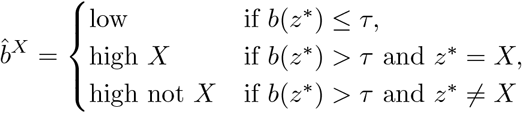

where *τ* is a threshold for partitioning. In our analyses, we set *τ* = 0.5.

With these partitions, we further define two binary variables along which to probe belief representation, confidence and preference, where:

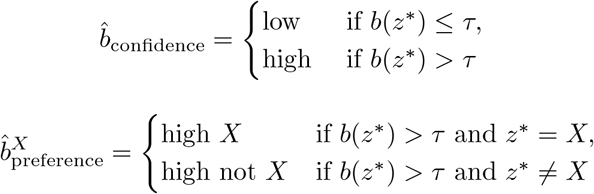

Single unit and decoding analyses of selected stimulus feature, reward, belief confidence, and belief preference (Figs 2, 3, 5, 6, 7, S3, S4, S5, S6) were all performed a feature at a time. For each feature, to guarantee that each recorded neuron corresponded to enough trials per-condition, we sub-selected sessions where the feature occurred as the hidden rule feature in at least 3 rule blocks throughout the session.

For analyses on pairs of features *X* and *Y*, we constrain our analyses to belief states that fall under the following three partitions:

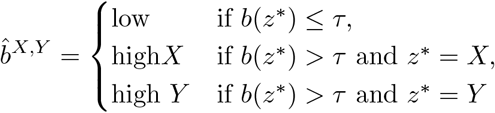

Population analyses of belief organization were performed across pairs of features (Figs 8, 9, S7, S8). To similarly guarantee enough trials per-condition, we only examined pairs of features where in at least 10 sessions, both features occurred as the hidden rule in at least 3 blocks each. Across both monkeys, out of the 66 possible feature pairs, 28 pairs satisfied this criteria, with 16 across feature dimension and 12 within feature dimension.

### 5.6 Single Unit Encoding

We are interested in whether single neuron firing rates encode relevant behavioral conditions of interest. To study this, we examine firing rates in trial periods aligned to stimulus onset and feedback onset, within a sliding 500ms window. Following notation from Kobak et al. [62], for a given neuron, in a 500ms window, the neuron’s firing rate on trial *k*, time bin *t*, and condition *c* can be denoted as *r*_*t,c,k*_. We can then compute a set of average firing rates over combinations of parameters, as:

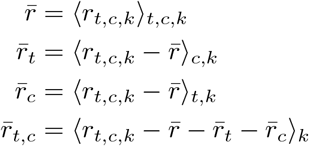

Here, 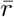 is the overall mean firing rate of the neuron in the window, 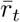 is the average time-varying firing rate with 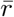 subtracted, 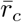 is average condition-varying firing rate with overall mean 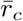 subtracted, and, 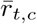 is the average time and condition varying firing rate with 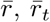, and 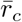 subtracted out. With these averages, the full firing rate *r*_*t,c,k*_ can be decomposed as:

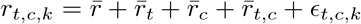

where *ϵ*_*t,c,k*_ is the residual noise term. The fraction of variance explained by condition and time is computed as:

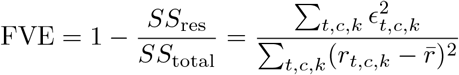

To test whether a neuron’s firing rate significantly encodes a condition in the time window, we compare the observed FVE to a null distribution generated by shuffling condition labels (see 5.10) and recomputing FVE. For *M* shuffles, the p-value is

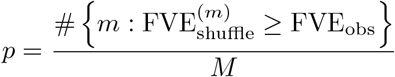

With *M* = 100. Neurons with *p* < 0.01 are considered to be significantly encoding the condition of interest at that time window.

### 5.7 Pseudo Population Construction

To explore the neural population representations of beliefs and other task variables, we combined neurons across recording sessions to create pseudo populations for decoding analyses. For some session *s* and some time bin of interest *t*, we record the firing rates of *N*_*s*_ neurons across *K*_*s*_ trials. The column vector of neural firing rates per trial *k* is denoted as 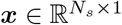. Correspondingly we denote the current binary condition label per trial *k* ∈ {1, …*K*_*s*_} as *y*(*k*) ∈ {*c*^+^, *c*^−^ }. Concatenating the labels and firing rates across trials gives us per-session, per time bin population firing rates and labels:

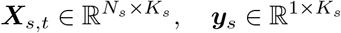

To construct our pseudo populations, for each recording session, we first group trials corresponding to each condition label *c* ∈ {*c*^+^, *c*^−^ }, and balance the number of trials in each condition by randomly sub-sampling trials for the condition with a larger number of trials, such that the same number of trials are present for each condition. For each condition, corresponding trials are then randomly split into train and test sets with a 80/20 ratio. For each session *s*, condition *c*, and time bin *t*, we then constructed pseudo-trials for training and testing by sampling with replacement the original trials corresponding to train/test sets. Drawing 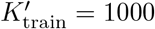 samples for training, and 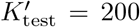 samples for testing, we form the corresponding neuronal firing rates into input data matrices:

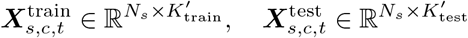

To construct pseudo populations, we concatenate our input data across all sessions, along the neuron dimension, such that pseudo trials are aligned, forming the training input matrix:

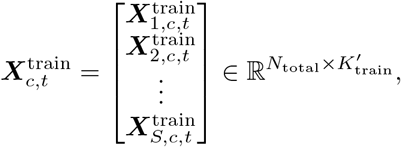

and the analogous input data matrix for testing data. Here, 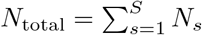. Training and testing data input matrices also correspond to labels vectors with the condition repeated:

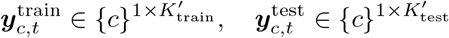

Finally, we concatenate our data across conditions, along the trial dimension, yielding the final training input data matrix:

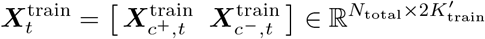

and training label vector:

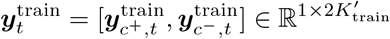

The input data matrix and label vector for testing data was analogously generated.

### 5.8 Population Decoders

For each task condition of interest, we trained regularized logistic regression models per time bin to decode task conditions from pseudo-population activity. Decoders were implemented as a single-layer linear classifier with dropout and batch normalization using the PyTorch library. For each pseudo-trial, the model for some some time bin *t* can be written as:

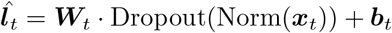

Where 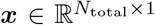 is the pseudo-trial’s firing rate inputs, 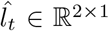 are the logits corresponding to *c*^+^ and *c*^−^ conditions, 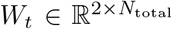 and *b* ∈ ℝ^2*×*1^ are learnable weights and biases, Norm(·) is batch normalization, and Dropout(·) randomly zeros out inputs during training at a rate of *p*_dropout_ = 0.5 to ensure regularization. Decoders are trained via cross-entropy loss on the training dataset.

During testing, predicted labels for each pseudo test trial *i* correspond to the class with the highest logit:

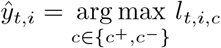

And test accuracy per-time bin can be assessed as:

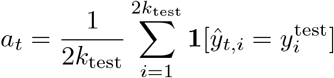

Decoders were evaluated a feature at a time, with recording sessions sub-selected to satisfied the criteria in 5.5. For each time bin, decoders were assessed across 8 different train/test splits The mean and S.E.M of these accuracies across all features and train/test splits are reported. p-values for each time bin’s accuracies are assessed via a permutation test against a shuffled null distribution (see 5.10). For each of our decoding analyses, we decoded from only units that were previously identified as significantly encoding the condition of interest in any time window of the trial (see 5.6). For preference and confidence decoding, to prevent the influence of behavioral confounds throughout the trial, we only examined trials where the feature of interest was selected and the trial was rewarded.

### 5.9 Cross Time Decoders

To assess decoding accuracy of task conditions across time bins, we take the decoders trained on some time bin *t*_train_ and evaluate them on neural activity recorded from time bin *t*_test_, we ensure the same train-test trial splits are preserved across time bins.

### 5.10 Generating Shuffle Distributions

One concern in our dataset might be that the some of the task conditions we are aiming to study can be slowly drifting in time, so units that are similarly slowly drifting in their firing rates may give rise to nonsense correlations [63]. To address this, in each of our analyses, we generated shuffled null distributions which best preserve the temporal structure in both neural activity and task conditions while breaking alignment across the two. For single unit encoding analyses in 5.6, we generated shuffles by circularly shifting the trial number for our task condition. Specifically, for a unit recorded on a session with *K*_*s*_ trials, let: 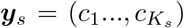 denote the sequence of original per-trial condition labels. We aim to generate a distribution of *M* shuffled labels that have been circularly shifted. For each shuffle *m* ∈ {1, …, *M* }, we shift our labels by a random number of trials *τ*_*m*_, sampled as:

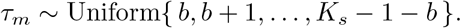

where *b* = 50 is the shift bounds to prevent a shift too close to the original trial. This generates a shuffled sequences of labels:

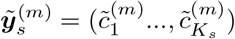

where for each trial *k*:

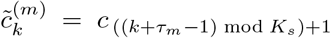

Firing rates for the single unit of interest is then associated with the shuffled labels, and we repeat our analysis for each of the *M* shuffles, generating our shuffled null distribution. For any population analyses, we take advantage of the fact that our analyses combines recordings across multiple sessions, and implement a session permutation shuffle [63]. To generate a shuffled dataset *m*, for a time point of interest *t*, for each session *s*, we correspond our neural activity ***X***_*s,t*_ with labels from another randomly chosen session *s*^*′*^ such that *s* ≠ *s*^*′*^. As the two sessions may contain different numbers of recorded trials, we compute the minimum shared trial length 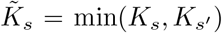 between the two, and clip both neural activity and labels to be 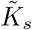 trials. Therefore, for shuffle *m*, session *s*, our shuffled dataset comprises of firing rates:

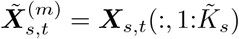

and labels:

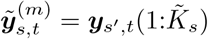

Across sessions, pseudo populations are generated in the same manner as in 5.7. For each of our analyses, shuffling *M* = 10 times provides us with a shuffled null distribution used to compare against true distributions.

### 5.11 Projecting Changes in Neural Activity

To study how belief-relevant neural activity changes across trials based off trial outcomes, we first let *c* ∈ ℂ denote the set of trial outcomes of interest. Outcomes can be possible combinations of prior belief partition, selected stimulus feature, or reward.

Similar to 5.7, for session *s* with *K* trials recorded, we generate per time bin *t* firing rate matrices and trial outcomes:

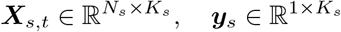

Where 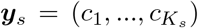 denotes the outcome labels per-trial. Here, trial outcomes can be possible combinations of stimuli selection, belief partition, and rewards. For each specific trial outcome of interest *c*, we are interested in the difference in neural activity between the trial where the outcome was observed, and the trial immediately after that. We find the set of all trials with outcome *c*:

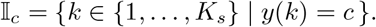

And generate a matrix differences in firing rates:

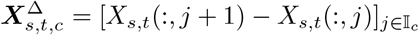

In the same manner as in 5.7, for each session, we draw *K*^*′*^ = 100 sample columns from our difference matrix, and concatenate data across sessions for form a combined matrix of:

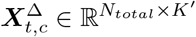

To project this neural activity onto corresponding preference and confidence decoder axes, we compute the relevant axes with decoder weights obtained in 5.13:

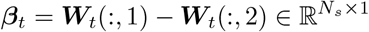

Across all pseudo-trials and time bins, the average projection onto the axis is computed as:

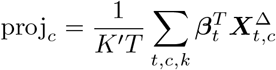

### 5.12 Cross Condition Generalization Performance

To assess the alignment of belief representations across features, we examine pairs of features at a time, sub-selecting for sessions which satisfy the criteria in 5.5. For a pair of features *X* and *Y*, we can separately define two sets of conditions:

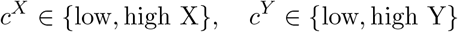

Population decoders are separately trained to classify either the *c*^*X*^ or *c*^*Y*^ conditions (5.13), and within-condition accuracy is assessed as test accuracy for each of the decoders. To assess alignment, we take decoders trained on *c*^*X*^ conditions, and evaluate how well they can classify *c*^*Y*^ conditions, and vice versa. To evaluate belief organization by dimension, we split pairs of features into within-feature-dimension and across-feature-dimension pairs:

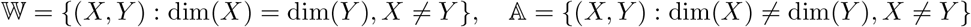

And evaluated whether the across condition accuracy differs between the two groups.

### 5.13 Belief Vector Similarities

To additionally assess the alignment of belief representations across features, we computed the cosine similarity across population vectors corresponding to increasing belief in features. Specifically, for some pair of features *X, Y*, on session *s*, and time bin *t*, we find population firing rates which correspond to each of the belief partitions of interest 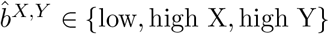 {low, high X, high Y}:

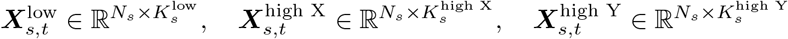

Similar to 5.7 and 5.11, we sample from *K*^*′*^ = 100 pseudo-trials from each of our population firing rates, and concatenate across all valid recording sessions to form pseudo populations:

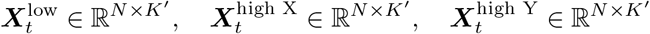

Per pseudo-trial *k*^*′*^, we define the belief vector for feature *X*, denoted as 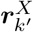 to be the population vector from low to high X partitions:

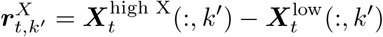

The average cosine similarity between belief vectors is then computed as:

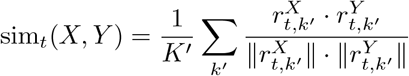

Correspondingly averaging across all time bins gives:

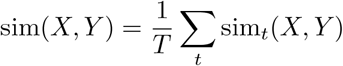

To evaluate the difference of belief vector similarities between within dimension feature pairs 𝕎 and across dimension feature pairs 𝔸, we define the average difference in within vs. across similarities as:

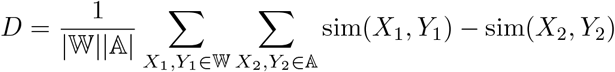

## 6 Acknowledgements

This work was supported by NSF GRFP 2023356971, NIH 5U19NS107609, NIH 1UF1NS126485, ORIP P51 OD010425, and the Simons Collaboration for the Global Brain.

We thank Charles Ian O’Leary, Sufia Ahmad, Megan Jutras, Kelly Morrisroe, and Sierra Schleufer for task design and data collection, and animal training. We additionally thank John Ferre, Po-Chen Kuo, W. Jeffery Johnston, Alison Duffy, David Bell, Scott Sterrett, and Sofia Landi for helpful discussions.

